# Regional macrophage specialization shapes intestinal neuroimmune function

**DOI:** 10.64898/2026.10.08.757647

**Authors:** Khaled Warasnhe, Juan Sebastian Ortiz Colmenares, Mario D’Ambrosio, Biagio Di Lorenzo, Yuanhang Liu, Sihan Ji, Magdalini Mischopoulou, Natalie R. Wertish, Mohamed H. Eldesouki, Xin-yi Chai, Jean X Jiang, Madhusudan Grover, Keith A. Sharkey, Arthur Beyder, Gianrico Farrugia, Gianluca Cipriani

**Affiliations:** Enteric Neuroscience Program (ENSP), Division of Gastroenterology & Hepatology, Department of Medicine, Mayo Clinic, Rochester, Minnesota; Department of Medicine and Surgery, LUM University, Casamassima, Italy; Department of Quantitative Health Sciences, Mayo Clinic, Rochester, Minnesota; Department of Neuroendocrine Pharmacology, School of Pharmacy, China Medical University, Shenyang, Liaoning Province, China; Department of Biochemistry and Structural Biology, University of Texas Health Science Center, San Antonio, Texas; Hotchkiss Brain Institute, Snyder Institute for Chronic Diseases and Department of Physiology & Pharmacology, Cumming School of Medicine, University of Calgary, Calgary, Alberta, Canada

## Abstract

Tissue-resident macrophages adapt to local environments to support organ-specific functions, but whether their regulation differs across regions of the same organ is poorly understood. The gastrointestinal tract provides an ideal system to address this question because its anatomically distinct regions perform specialized physiological functions and contain different cellular environments. Here, we found that intestinal muscularis macrophages (MMs) exhibited distinct regional trophic requirements and neuroimmune functions across the gut. Small-intestinal MMs were highly sensitive to CSF1 neutralization but recovered robustly, whereas colonic MMs were less sensitive to CSF1 depletion yet showed limited recovery. Disruption of CSF1-dependent MMs selectively reduced nitrergic neurons and altered motility in the small intestine, whereas the colonic neuronal compartment was comparatively preserved. *Csf1* was enriched in small-intestinal nNOS^+^ neurons, and neuron-specific *Csf1* deletion reduced both MMs and nitrergic neurons and altered small-intestinal transit, with limited effects in the colon. Together, these findings reveal that macrophage trophic requirements and neuroimmune functions are regionally specialized within the intestine and identify neuronal CSF1 as a key component of the trophic relationship between nitrergic neurons and MMs in the small intestine.

## INTRODUCTION

Tissue-resident macrophages acquire specialized identities and functions in response to signals from the tissues they inhabit (1). This adaptation allows macrophages to support organ-specific processes ranging from tissue remodeling and metabolism to neuronal and vascular homeostasis. However, organs themselves are spatially heterogeneous, containing anatomically and functionally distinct regions with different cellular environments and physiological demands. It remains unclear whether macrophages residing in different regions of the same organ have distinct trophic requirements for their survival and maintenance and whether they establish functional interactions specific to their local environment. Defining this level of specialization is important for understanding how local tissue environments organize macrophage function in both homeostasis and disease.

The gastrointestinal (GI) tract provides an ideal system for addressing this question, because each of its regions carries out distinct physiological functions and harbors its own specialized immune and neural environments. Intestinal macrophages occupy multiple anatomical compartments and contribute to epithelial, immune, neuronal, and motor homeostasis (2, 3). Within the muscularis externa, muscularis macrophages (MMs) reside in close association with enteric neurons, glia, interstitial cells of Cajal (ICCs), and stromal populations, where they contribute to enteric neuronal homeostasis and GI motility. Communication between MMs and surrounding cells relies on several signaling pathways, including colony-stimulating factor 1 (CSF1), a major trophic factor for tissue macrophages. Enteric neurons provide CSF1 to neighboring CSF1R^+^MMs (4), while additional cellular populations contribute to local CSF1 availability (5). Yet MMs persist even when CSF1-CSF1R signaling is disrupted (6, 7), suggesting that macrophage trophic requirements may not be uniform across the gut but instead vary with regional specialization of the local neuroimmune environment.

Here, we show that MMs residing in different intestinal regions exhibit distinct trophic requirements, responses to perturbation, and functional relationships within the enteric nervous system. Small-intestinal and colonic MMs differed markedly in their sensitivity to CSF1 neutralization and, unexpectedly, resistance to depletion did not predict subsequent recovery. These regional differences extended beyond survival to include macrophage composition, transcriptional signature, and communication with surrounding cells. Functionally, disruption of CSF1-dependent MMs preferentially altered small-intestinal nitrergic neurons and motility. Conversely, *Csf1* was enriched in nNOS^+^ enteric neurons, and neuronal *Csf1* deletion selectively reduced small-intestinal MMs and nitrergic neurons. Together, these findings establish regional specialization as a fundamental feature of intestinal macrophages, showing that MMs form distinct trophic and neuroimmune relationships across functionally different regions of the gut.

## RESULTS

### Regional CSF1 dependence defines distinct muscularis macrophage populations

We started by exploring CSF1, an established major trophic factor for macrophages, which supports the maintenance and survival of tissue-resident macrophages (7). While parts of the same organ, small intestine (SI) and colon have distinct functions and neuro-immune repertoires. We therefore asked whether MMs depend on CSF1 uniformly across the GI tract. We administered αCSF1 to adult mice and quantified MHCII⁺ MMs by whole-mount immunohistochemistry (**Figures 1a,c**). Seven days of αCSF1 treatment markedly reduced MM abundance in the stomach, SI, and colon without significantly alter intestinal length or body weight during the treatment period (**Supplementary Figure 1a-c**). Interestingly, the extent of depletion differed substantially across gut regions. MMs were mostly depleted in the stomach and SI, whereas a substantially larger macrophage population persisted in the colon (**Figures 1b,c**).

**Figure 1.**
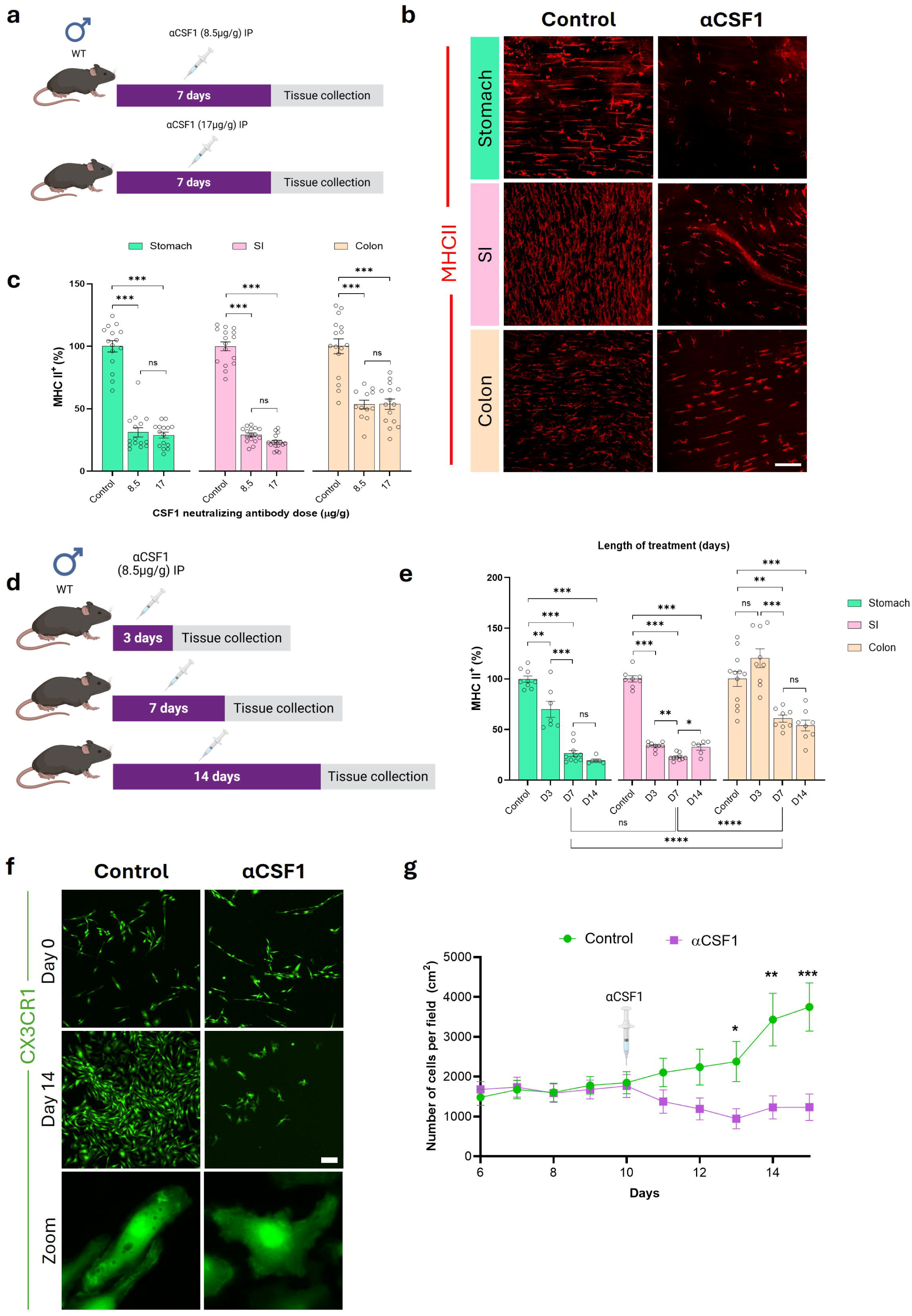
Regional CSF1 dependence defines distinct muscularis macrophage populations. (a) Schematic representation of αCSF1 treatment paradigms used to assess dose-dependent depletion of MMs in the stomach, SI, and colon. (b) Representative whole-mount images of MHCII⁺ MMs in the stomach, SI, and colon from control and αCSF1-treated mice. (c) Quantification of MHCII⁺ MMs in the stomach, SI, and colon following αCSF1 treatment at the indicated doses. (d) Schematic representation of the αCSF1 time-course experiment. Mice received αCSF1 for 3, 7, or 14 days before tissue collection. (e) Quantification of MHCII⁺ MMs in the stomach, SI, and colon across the αCSF1 treatment time course. (f) Representative images of FACS-isolated CX3CR1-GFP⁺ MMs cultured ex vivo under control or αCSF1 conditions. (g) Quantification of GFP⁺ macrophage abundance during ex vivo culture under control and αCSF1 conditions. Bar graphs represent the mean ± SEM; ∗ *P* < 0.05, ∗∗ *P* < 0.01, ∗∗∗ *P* < 0.001, ns, nonsignificant (one-way ANOVA with Bonferroni’s multiple comparison). Scale bar: 100 μm.

We next asked whether residual MMs reflected incomplete CSF1 neutralization. Increasing the αCSF1 dose from 8.5 to 17 μg/g did not produce further depletion in any GI region (**Figure 1a,c).** Similarly, extending treatment revealed distinct regional depletion kinetics followed by a plateau despite continued CSF1 neutralization (**Figure 1d,e**). In the stomach and SI, MM abundance declined significantly by day 3, decreased further by day 7, and showed no significant additional depletion by day 14. In contrast, the decline in colonic MM was less than in the other regions of the gut, becoming significant by day 7, with no further significant reduction by day 14. RT-qPCR revealed that *Csf1* expression was significantly higher in the SI than in the colon, whereas *Csf1r* expression was comparable between the two regions (**Supplementary Figure 1d**). Thus, the greater sensitivity of SI MMs to CSF1 neutralization occurred in a tissue environment characterized by higher *Csf1* expression, implicating regional differences in CSF1-dependent trophic organization.

To determine whether this differential sensitivity to CSF1 neutralization was maintained outside the intestinal tissue environment, we isolated muscularis externa CX3CR1-GFP⁺ MMs which are CSF1R-dependent (4) by FACS (**Supplementary Figures 1e,f**) and examined their CSF1 dependency *in vitro*. Control macrophages progressively expanded, whereas αCSF1 treatment prevented this expansion and reduced macrophage number (**Figure 1f,g**). Nevertheless, a subset of GFP⁺ macrophages persisted despite sustained CSF1 neutralization and acquired a less elongated, more rounded morphology (**Figure 1f**). Thus, heterogeneous sensitivity to CSF1 neutralization was recapitulated *in vitro*. These *in vitro* findings indicate that heterogeneous sensitivity to CSF1 neutralization can persist outside the intact tissue environment.

Together, these findings identify regionally distinct trophic requirements among intestinal MMs. Stomach and SI MMs exhibit high sensitivity to CSF1 neutralization, whereas the colon retains a substantially larger macrophage population despite increased αCSF1 dose and prolonged treatment. The persistence of residual MMs during sustained CSF1 neutralization, both in vivo and *in vitro*, supports the presence of macrophage populations with differential dependence on CSF1-mediated trophic support. These findings suggest that regional CSF1 dependence is an important determinant of intestinal MM heterogeneity and prompted us to examine whether sensitivity to CSF1 neutralization predicts subsequent macrophage recovery.

### Transient CSF1 depletion reveals region-specific muscularis macrophage recovery

Having established that intestinal MMs differ in their sensitivity to CSF1 neutralization, we next asked whether these regional differences were also reflected in their response after CSF1 availability was restored. Mice were treated with αCSF1 for 7 days, after which treatment was discontinued, and MMs abundance and phenotype were assessed following a 6-week recovery period (**Figure 2a**). MM recovery differed markedly across gut regions. In the stomach and SI, MM numbers returned to levels comparable to untreated controls, whereas colonic MMs showed no recovery (**Figure 2a,b**). In both regions, repopulating MMs returned to their normal distribution throughout the muscularis externa and re-established close spatial association with myenteric neurons, despite the incomplete numerical recovery seen in the colon (**Figure 2c**). Thus, intestinal MM niches exhibit distinct regional capacities for repopulation following transient CSF1 depletion.

**Figure 2.**
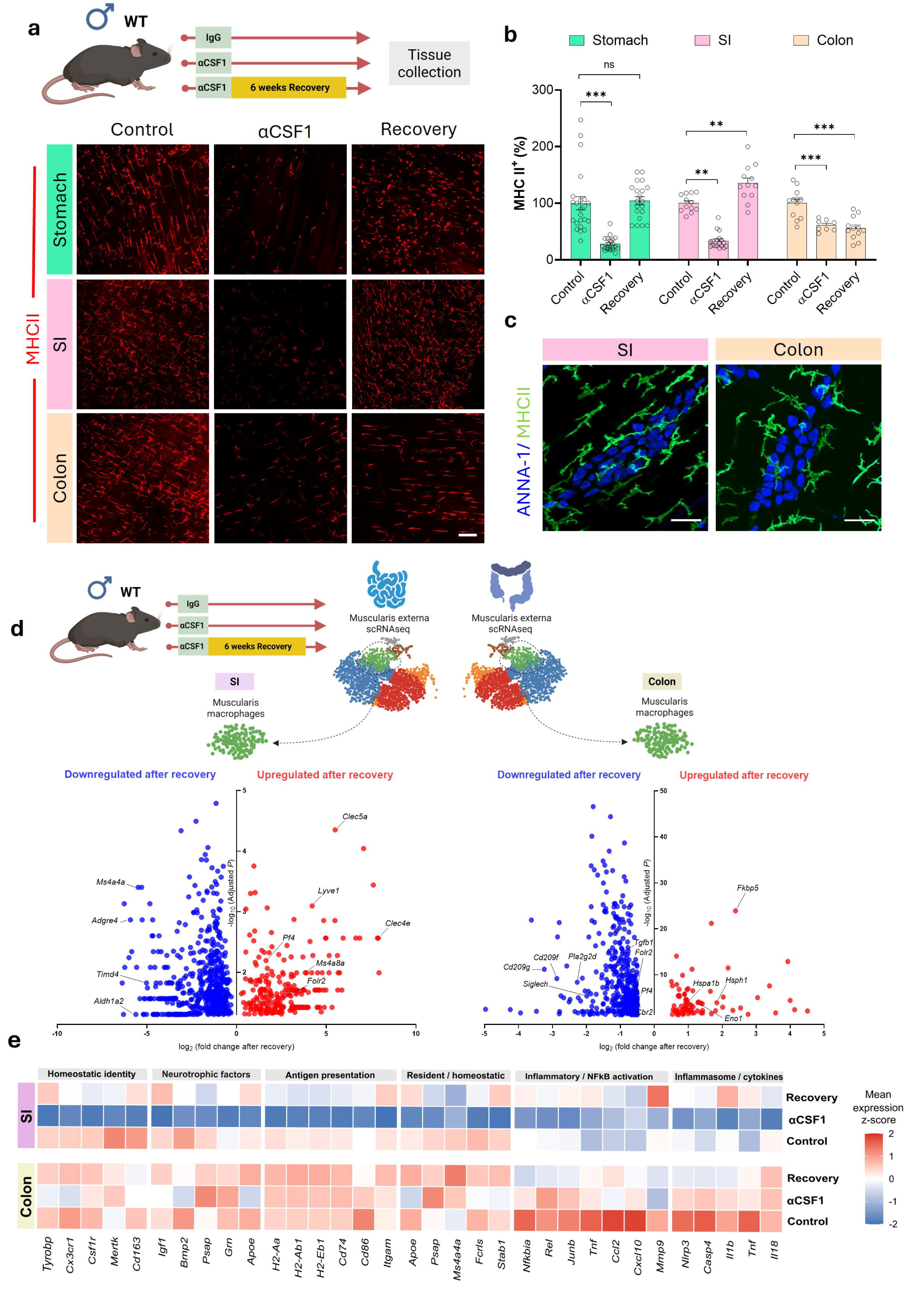
Transient CSF1 depletion reveals region-specific muscularis macrophage recovery. Mice received αCSF1 for 7 days followed by a 6-week recovery period without further αCSF1 treatment. (a) Representative whole-mount images of MHCII⁺ MMs in the stomach, SI, and colon across control, αCSF1-treated, and recovery conditions. (b) Quantification of MHCII⁺ MMs in the stomach, SI, and colon in control, αCSF1-treated, and recovered mice. (c) Representative images of MMs in the SI and colon following the 6-week recovery period. (d) Volcano plot of selected differentially expressed macrophage genes across control, αCSF1-treated, and recovery conditions in the SI and colon. (e) Macrophage transcriptional program scores across control, αCSF1-treated, and recovery conditions in the SI and colon, including homeostatic, neurotrophic, antigen-presentation, resident/tissue-adaptation, and inflammatory programs. Each point represents one image (*n =* 4-5 mice per group). Bar graphs represent the mean ± SEM; ∗∗ *P* < 0.01, ∗∗∗ *P* < 0.001, ns, nonsignificant (one-way ANOVA with Bonferroni’s multiple comparison). Scale bar: (a) 100 μm, (c) 50 μm.

We next wondered about the identities of the restored macrophages in the SI and colon. Single-cell RNA sequencing of the muscularis externa revealed persistent transcriptional remodeling after recovery, with marked differences between the SI and colon (**Figure 2d**). Despite robust numerical repopulation in SI, recovered MMs did not fully re-acquire their steady-state transcriptional signature (**Figure 2d,e**). Recovered MMs populations in the SI exhibited a mixed transcriptional phenotype, with increased expression of tissue-resident/tissue-supportive genes (*Lyve1*, *Folr2*), persistent reduction of other steady-state resident-associated genes (*Timd4*, *Adgre4*, *Aldh1a2*), and increased expression of additional state-associated genes (*Clec5a*, *Clec4a*, *Pf4*) (**Figure 2d**) (8). Analysis of broader transcriptional programs across control, αCSF1-treated, and recovery conditions further revealed remodeling of homeostatic, neurotrophic, antigen-presentation, tissue-adaptation, and inflammatory programs (**Figure 2e**; **Supplementary Figure 2a**). In the SI, CSF1 depletion broadly suppressed homeostatic, neurotrophic, antigen-presentation, and tissue-adaptation programs in MMs. During recovery, many of these programs returned toward control levels, whereas others remained only partially restored. Thus, in the SI, MMs number restoration was not accompanied by a fully restored macrophage identity. Colonic MMs followed a distinct trajectory. Consistent with incomplete numerical recovery, recovered colonic MMs remained transcriptionally remodeled, showing persistent alterations in resident/tissue-adaptation programs and increased expression of stress-associated genes, including *Fkbp5*, *Hspa1a*, and *Hsph1* (**Figure 2d,e**; **Supplementary Figure 2a**). SI and colon therefore differed not only in the extent of macrophage repopulation but also in the molecular signature of the reconstructed MMs.

These divergent outcomes were not explained by persistent differences in tissue CSF1 abundance, as *Csf1* expression was comparable among control, treated, and recovered tissues in both the SI and colon (**Supplementary Figure 2b**). Moreover, CX3CR1-GFP⁺ MMs surviving CSF1 neutralization *in vitro* resumed expansion after antibody withdrawal, although their numbers remained lower than those of control-treated MMs (**Supplementary Figure 2c,d**). Thus, macrophages that persist during transient CSF1 neutralization retain the capacity to resume expansion following restoration of CSF1 availability *in vitro*.

Together, these findings demonstrate that intestinal MMs populations differ not only in their sensitivity to CSF1 neutralization but also in their response after the perturbation is removed. The highly CSF1-sensitive SI population underwent robust numerical recovery, whereas the comparatively preserved colonic population remained numerically reduced. Moreover, restoration of MMs abundance in the SI was not accompanied by complete restoration of the steady-state transcriptional state. Thus, trophic sensitivity, numerical recovery, and molecular restoration represent distinct properties of regionally specialized intestinal MMs populations.

### Macrophage recovery follows distinct cellular trajectories across intestinal regions

Having observed region-specific differences in MMs recovery, we next asked to what extent macrophages present before CSF1 neutralization contribute to the MMs population after the recovery period. To track MMs present before depletion, *LysM*^CreERT2^xAi9 mice received tamoxifen before αCSF1 treatment, followed by a 6-week recovery period (**Figure 3a**). Tamoxifen induced robust tdTomato labeling of *LysM*-expressing cells, which was confirmed by flow cytometry (**Supplementary Figure 3a,b**).

**Figure 3.**
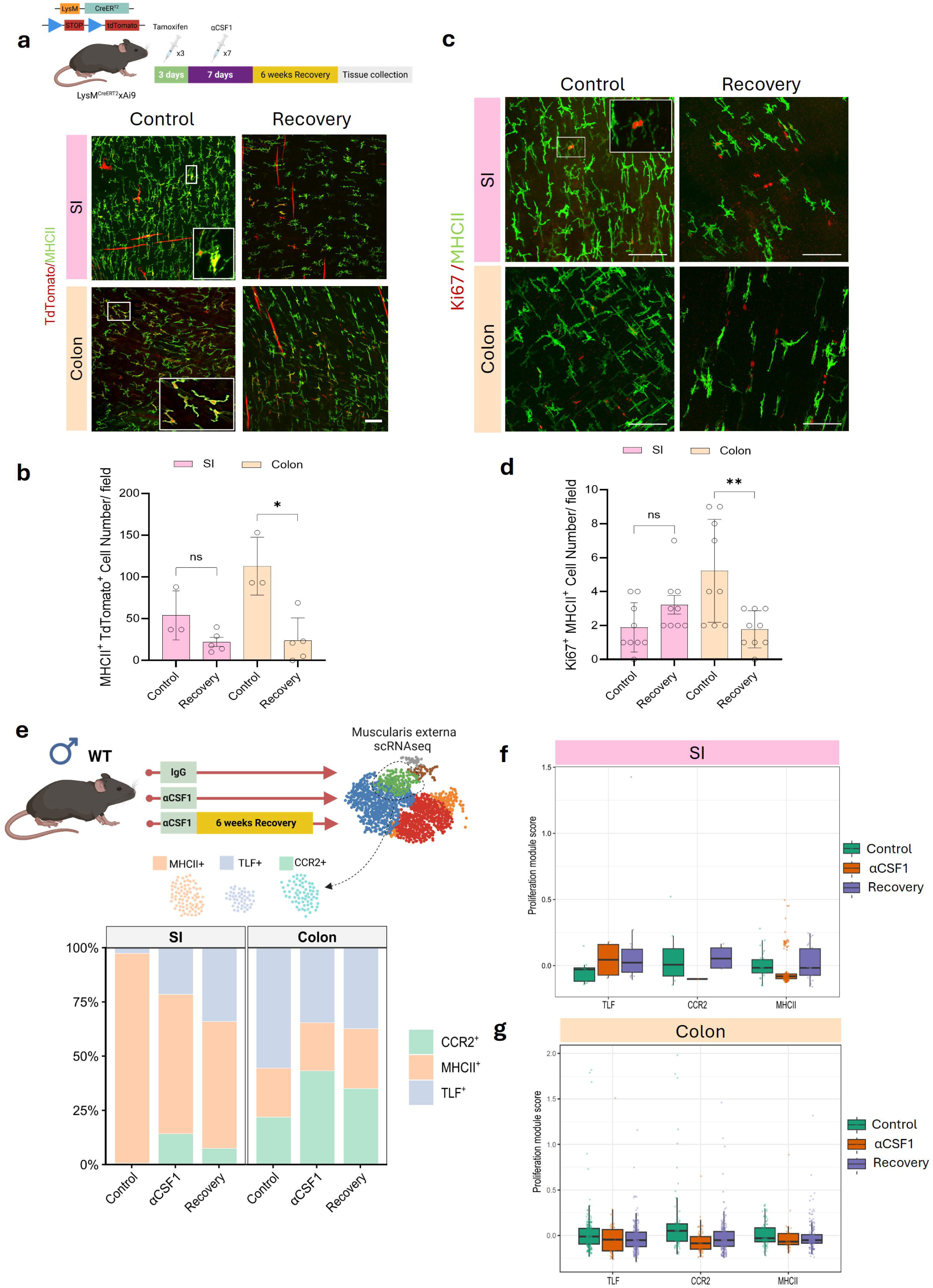
Macrophage recovery follows distinct cellular trajectories across intestinal regions. (a) Schematic of the inducible lineage-labeling and recovery paradigm in *LysM* ^CreERT2^xAi9 mice and representative whole-mount images of tdTomato⁺MHCII⁺ macrophages in the SI and colon from control and recovered mice. Cells were labeled before αCSF1 treatment and analyzed following a 6-week recovery period. (b) Quantification of tdTomato⁺MHCII⁺ macrophages in the SI and colon from control and recovered mice. (c) Representative whole-mount images of Ki67⁺MHCII⁺ macrophages in the SI and colon from control and recovered mice. (d) Quantification of Ki67⁺MHCII⁺ macrophages in the SI and colon following recovery (*n =* 3). (e) Relative abundance of TLF-, MHCII-, and CCR2-associated macrophage populations across control, αCSF1-treated, and recovery conditions in the SI and colon. (f) Proliferation module score across TLF-, CCR2-, and MHCII-associated macrophage populations in SI. (g) Proliferation module score across the corresponding macrophage populations in the colon. Each point represents one biological replicate unless otherwise indicated. Bar graphs represent the mean ± SEM; ∗ *P* < 0.05, ∗∗ *P* < 0.01 ns, nonsignificant (unpaired *t*-test). Scale bar: 50 μm.

Lineage labeling revealed marked regional differences in the contribution of macrophages labeled before depletion to the restored MMs pool. In the SI, tdTomato⁺ MHCII⁺ macrophages remained abundant after recovery and were not significantly reduced relative to controls. In contrast, tdTomato⁺ MHCII⁺ macrophages were markedly reduced in the recovered colon (**Figure 3a,b**). Thus, macrophages labeled before CSF1 depletion remained a substantial component of the recovered SI macrophage pool, whereas their contribution to the recovered colonic compartment was markedly reduced. Because local proliferation contributes to the maintenance of tissue-resident macrophage populations (1), we next examined MM proliferative activity following recovery. Ki67⁺ MHCII⁺ MMs were maintained in the SI following recovery but were significantly reduced in the recovered colon (**Figure 3c,d**), in line with a reduction of pre-labeled tissue resident MMs. Thus, persistence of the pre-depletion labeled compartment in the SI was accompanied by preserved proliferative activity, whereas both were diminished during colonic MM recovery.

Single-cell RNA sequencing further revealed region-specific changes in MMs composition during depletion and recovery (**Figure 3e**; **Supplementary Figure 3c,d**). MMs were classified into 3 subpopulations based on TLF-associated genes (*Timd4*, *Lyve1*, and *Folr2*), MHCII-associated genes (*Cd52*, *H2-Eb1*, *H2-Aa*, and *H2-Ab1*), and the recruitment-associated marker *Ccr2* as previously described (9). These populations represent transcriptionally and functionally distinct macrophage states with different relationships to tissue residency, local maintenance, and monocyte recruitment (ref). TLF-associated macrophages are enriched for genes characteristic of long-lived, tissue-adapted macrophages, whereas CCR2-associated macrophages are associated with monocyte recruitment. MHCII-associated macrophages are distinguished by enrichment of antigen-presentation programs. These subpopulations differed in abundance between the steady-state SI and colon and followed distinct trajectories during depletion and recovery. In the SI, CSF1 depletion reduced the dominant MHCII^+^-associated population and markedly increased the relative abundance of CCR2⁺-associated and TLF^+^-associated macrophages. During recovery, CCR2⁺-associated population declined, whereas the TLF^+^-associated population increased and together with the MHCII+-associated population, constituted the majority of the recovered MMs pool (**Figure 3e**). Thus, SI macrophage composition shifted from increased representation of recruitment-associated macrophages during depletion toward a composition more closely resembling the steady-state compartment during recovery, although complete restoration was not achieved. Proliferation module scores were increased in TLF-associated macrophages during CSF1 depletion (**Figure 3f**), indicating enrichment of a proliferation-associated transcriptional signature within this population.

The colon followed a distinct trajectory. The steady-state colon contained substantial TLF^+^, MHCII^+^, and CCR2^+^-associated subpopulations, and CSF1 depletion further increased the relative representation of CCR2⁺-associated macrophages, as observed in the SI. However, the TLF^+^-associated population was reduced during depletion and never restored following recovery, consistent with the absence of an increase in the TLF proliferation module score (**Figure 3e**). Recruitment module scores further revealed persistent region- and state-dependent remodeling that was not fully captured by CCR2^+^-associated population frequencies alone (**Supplementary Figure 3e,f**).

Together, lineage labeling, proliferation, and single-cell analyses reveal distinct cellular trajectories of MMs recovery across the gut regions. SI recovery is associated with persistence of macrophages labeled before depletion, preserved proliferative activity, and substantial compositional restoration. In contrast, colonic recovery is characterized by reduced representation of macrophages labeled before depletion, diminished proliferative activity, and persistent recruitment-associated remodeling. These divergent cellular trajectories provide a potential cellular basis for robust numerical repopulation in SI and incomplete recovery of the colonic MM compartment.

### Macrophage recovery fails to restore communication with neurons in the small intestine

Because macrophages following restoration remained transcriptionally distinct from their steady-state counterparts, we asked whether numerical repopulation was accompanied by restoration of macrophage communication with surrounding cells. Single-cell RNA sequencing identified the major cellular populations of the SI and colon, including macrophages, enteric neurons, glia, ICCs, fibroblasts, endothelial cells, pericytes, and smooth muscle cells (**Figure 4a**). We therefore performed ligand-receptor analysis to infer macrophage-centered communication networks across control, depletion, and recovery conditions, focusing on cellular populations associated with the myenteric plexus niche (**Figure 4b**).

**Figure 4.**
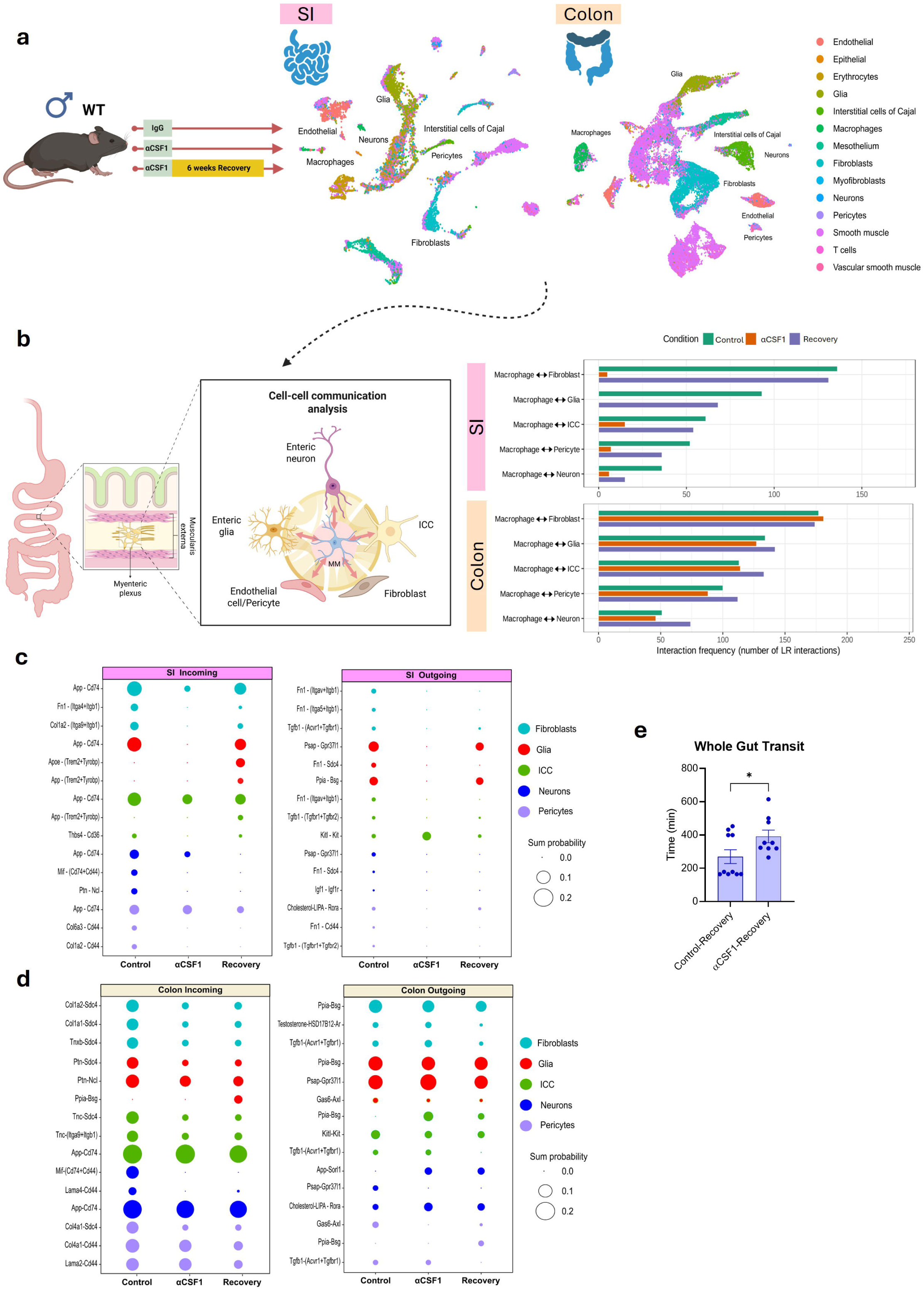
Macrophage recovery fails to restore communication with neurons in the small intestine. (a) UMAP visualization of single-cell RNA-seq data from the SI and colonic muscularis externa showing major cellular populations, including macrophages, enteric neurons and glia, interstitial cells of Cajal, fibroblasts, endothelial cells, pericytes, and smooth muscle cells. (b) Schematic representation of the myenteric plexus niche and predicted macrophage-centered communication with associated cellular populations, together with interaction frequency across control, αCSF1-treated, and recovery conditions in the SI and colon. (c) Selected predicted incoming and outgoing ligand-receptor interactions of SI macrophages across control, αCSF1-treated, and recovery conditions. Dot size represents summed inferred communication probability. (d) Selected predicted incoming and outgoing ligand-receptor interactions of colonic macrophages across control, αCSF1-treated, and recovery conditions. (e) Whole-gut transit time measured 6 weeks after cessation of αCSF1 treatment in control- and αCSF1-recovery mice.

CSF1 neutralization markedly disrupted predicted macrophage communication in the SI, particularly with fibroblasts, glia, ICCs, and enteric neurons (**Figure 4b**). Following 6 weeks of recovery, interactions with several niche populations were re-established, whereas communication with enteric neurons and glia remained partially altered. The extent of recovery varied among cell types, indicating selective rather than uniform restoration of macrophage signaling relationships. These data further highlight the broad integration of CSF1-dependent MMs with multiple cellular components of the neuromuscular compartment. Thus, despite robust numerical repopulation, SI MMs failed to fully re-establish their steady-state communication with enteric neurons and glia.

The colon followed a distinct trajectory. Predicted macrophage communication was comparatively preserved during CSF1 neutralization, consistent with greater persistence of colonic MMs, but interactions with glia, ICCs, pericytes, and neurons remained altered after recovery (**Figure 4b**). Broader analysis of macrophage interactions across muscularis externa populations similarly revealed distinct regional trajectories during depletion and recovery (**Supplementary Figure 4a**). Thus, the SI and colon differed both in the magnitude of communication disruption and in the pattern of communication observed in the recovery.

Analysis of individual incoming and outgoing ligand-receptor interactions further revealed region-specific remodeling of macrophage signaling (**Figures 4c,d**). In the SI, multiple predicted interactions with fibroblasts, glia, ICCs, neurons, and pericytes were lost or reduced during CSF1 neutralization. Some re-emerged during recovery, whereas others remained altered or acquired recovery-specific patterns. Colonic macrophages exhibited a distinct pattern, with several interactions comparatively preserved during CSF1 neutralization and recovery. Thus, macrophage repopulation in the SI did not fully restore the frequency of the interactions in the SI, and the trajectory of communication remodeling differed between the SI and colon.

Consistent with the broader regional differences in macrophage communication, *Csf1* expression was reduced across multiple SI cell populations following αCSF1 treatment, whereas no comparable change was observed in the colon (**Supplementary Figures 4b,c**). Immunohistochemical imaging further showed that residual MMs following CSF1 neutralization were preferentially associated with PECAM1⁺ blood vessels (**Supplementary Figure 4d**). These findings indicate that CSF1 neutralization remodels not only macrophage communication but also the spatial organization of the residual macrophage compartment, raising the possibility that distinct microanatomical environments contribute to macrophage persistence during CSF1 neutralization. We next asked whether incomplete restoration of macrophage communication was accompanied by persistent alterations in GI motility. Six weeks after cessation of αCSF1 treatment, mice exhibited significantly delayed whole-gut transit compared with recovery controls (**Figure 4e**). Importantly, this dysfunction persisted despite robust numerical repopulation of SI macrophages.

These data demonstrate that transient CSF1 disruption produces persistent remodeling of macrophage communication within the SI neuromuscular niche, with comparatively limited effects in the colon. Despite complete numerical restoration of SI MMs following recovery, predicted communication with enteric neurons and glia remained impaired. These persistent alterations in macrophage neuromuscular communication were accompanied by sustained changes in GI motility.

### CSF1-dependent macrophages selectively support small intestinal nitrergic neurons

Given the close association between MMs and the enteric nervous system (2), and the reduction of interaction frequency between MMs and enteric neurons following perturbation, we asked whether loss of CSF1-dependent MMs altered enteric neuronal organization in a region-dependent manner. Whole-mount preparations from the stomach, SI, and colon were immunolabeled for the pan-neuronal marker ANNA-1 and neuronal nitric oxide synthase (nNOS^+^) following 7 days of αCSF1 treatment (**Figure 5a**). Total ANNA-1⁺ neuronal number was unchanged in the stomach and colon but significantly reduced in the SI (**Figure 5a,b**). nNOS⁺ neurons were similarly reduced selectively in the SI, with no significant change in the stomach or colon (**Figure 5a,c**). Accordingly, the ratio of nNOS⁺ neurons to total ANNA-1⁺ neurons was reduced in the SI but remained unchanged in the other regions (**Figure 5d**). Thus, disruption of the CSF1-dependent macrophage niche preferentially affected the SI nitrergic neuronal compartment.

**Figure 5.**
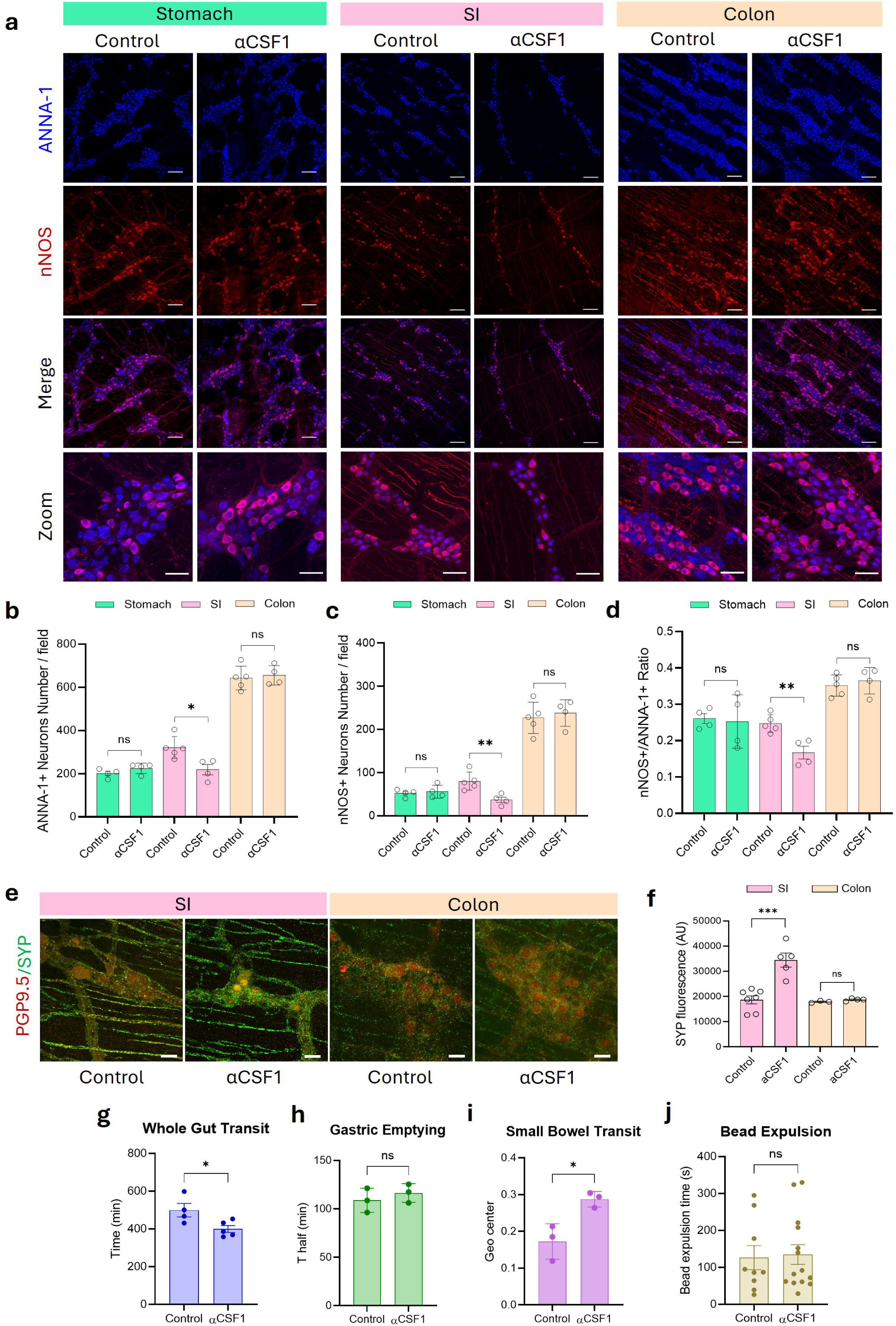
CSF1-dependent macrophages selectively support small intestinal nitrergic neurons. (a) Representative whole-mount images of ANNA-1⁺ and nNOS⁺ enteric neurons in the stomach, SI, and colon from control and αCSF1-treated mice for 7 days. (b) Quantification of total ANNA-1⁺ enteric neurons in the stomach, SI, and colon following αCSF1 treatment. (c) Quantification of nNOS⁺ enteric neurons in the stomach, SI, and colon following αCSF1 treatment. (d) Proportion of nNOS⁺ neurons to total ANNA-1⁺ neurons in the stomach, SI, and colon. (e) Representative synaptophysin immunolabeling within the myenteric plexus of the SI and colon from control and αCSF1-treated mice. (f) Quantification of synaptophysin immunoreactivity in the SI and colon. (g) Whole-gut transit time in control and αCSF1-treated mice. (h) Gastric emptying in control and αCSF1-treated mice. (i) Small bowel transit in control and αCSF1-treated mice. (j) Distal colonic bead-expulsion time in control and αCSF1-treated mice. Each point represents one biological replicate unless otherwise indicated. Bar graphs represent the mean ± SEM; ∗ *P* < 0.05, ∗∗ *P* < 0.01, ∗∗∗ *P* < 0.001, ns, nonsignificant (unpaired *t*-test). Scale bar: 100 μm (a), 50 μm (e).

To determine whether this remodeling extended across other enteric neuronal subtypes, we examined cholinergic neurons using ChATxAi3 reporter mice (**Supplementary Figure 5a**). In contrast to nNOS⁺ neurons, ChAT⁺ neuronal number was unchanged following αCSF1 treatment in the stomach, SI, and colon (**Supplementary Figure 5b,c**). Total ANNA-1⁺ neurons were again reduced in SI, whereas the relative representation of ChAT⁺ neurons was preserved. Thus, CSF1 neutralization does not induce uniform enteric neuronal loss but selectively remodels the SI nitrergic neuronal compartment.

We next assessed whether this neuronal remodeling was accompanied by changes in synaptic organization. Synaptophysin immunoreactivity within the myenteric plexus was significantly increased following αCSF1 treatment in the SI but remained unchanged in the colon (**Figure 5e,f**). These findings indicate that disruption of CSF1-dependent MMs is accompanied by remodeling of both nitrergic neuronal composition and synaptic architecture specifically within the SI.

We next examined whether these neural changes were associated with regional alterations in GI motility. αCSF1 treatment significantly shortened whole-gut transit time (**Figure 5g**) and increased small-bowel transit (**Figure 5i**), whereas gastric emptying (**Figure 5h**) and distal colonic bead expulsion (**Figure 5j**) were unchanged. Thus, the regional pattern of motility dysfunction paralleled the neuronal phenotype, with remodeling of the SI enteric nervous system accompanied by altered SI transit while colonic neuronal composition and basal colonic motility remained preserved.

To determine whether neuronal remodeling depended on the extent of macrophage depletion, we independently targeted CSF1R, producing near-complete depletion of MMs in both the SI and colon (**Supplementary Figure 6a**), as previously described (4). αCSF1R treatment accelerated whole-gut transit and distal colonic bead expulsion, whereas gastric emptying remained unchanged and small-bowel transit showed a trend toward acceleration (**Supplementary Figures 6b-e**). Near-complete macrophage depletion was also accompanied by reductions in total ANNA-1⁺ and nNOS⁺ neuronal numbers in both the SI and colon (**Supplementary Figures 6f-h**). However, the proportion of nNOS⁺ neurons among total ANNA-1⁺ neurons remained unchanged in both regions (**Supplementary Figure 6i**). Thus, extensive CSF1R-mediated macrophage depletion produced broader changes in neuronal composition without the selective regional alteration in nitrergic neurons observed following CSF1 neutralization.

Together, these findings identify a region- and neuronal subtype-specific relationship between CSF1-dependent MMs and the enteric nervous system. CSF1 neutralization selectively reduces SI nitrergic neurons while sparing cholinergic neurons, increases synaptophysin immunoreactivity within the SI myenteric plexus, and alters small-intestinal transit. In contrast, near-complete αCSF1R-mediated macrophage depletion produces broader reductions in neuronal abundance without selectively altering regional nitrergic representation. These findings indicate that enteric neuronal remodeling does not simply rely on the extent of macrophage depletion but differs according to the nature and regional context of macrophage niche disruption.

### Nitrergic neurons provide CSF1-dependent trophic support to small-intestinal macrophages

The selective vulnerability of SI nitrergic neurons following CSF1-dependent macrophage depletion prompted us to ask whether this neuronal subtype also contributes to the trophic environment that maintains MMs. Analysis of single-cell transcriptomic datasets revealed marked enrichment of *Csf1* expression in nNOS^+^ enteric neurons in the SI, whereas ChAT⁺ neurons showed little or no *Csf1* expression (**Figure 6a**). Neuronal *Csf1* expression was substantially lower in the colon, revealing both regional and neuronal subtype specificity in the neuronal CSF1 environment.

**Figure 6.**
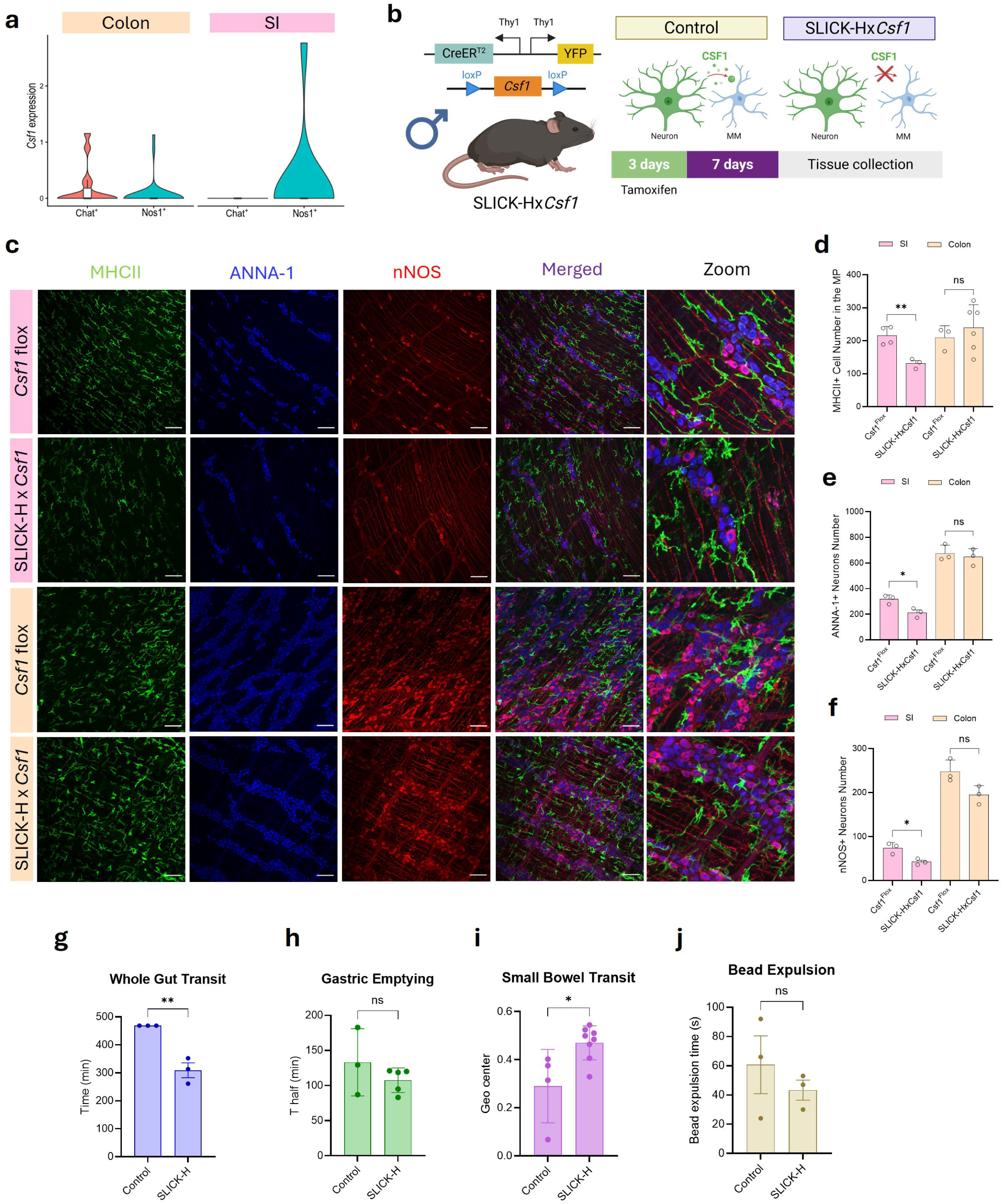
Nitrergic neurons provide CSF1-dependent trophic support to small-intestinal macrophages. (a) Single-cell transcriptomic analysis of neuronal *Csf1* expression in ChAT⁺ and nNOS⁺ enteric neuronal populations in the SI and colon. (b) Schematic representation of tamoxifen-induced neuronal *Csf1* deletion in SLICK-Hx*Csf1* mice and the proposed effect of neuronal CSF1 loss on the MMs niche. (c) Representative whole-mount images of MHCII⁺ MMs, ANNA-1⁺ enteric neurons, and nNOS⁺ neurons in the SI and colon from control and SLICK-Hx*Csf1* mice. (d) Quantification of MHCII⁺ MMs in the myenteric plexus region of the SI and colon. (e) Quantification of ANNA-1⁺ enteric neurons in the SI and colon following neuronal Csf1 deletion. (f) Quantification of nNOS⁺ enteric neurons. (g) Whole-gut transit time. (h) Gastric emptying. (i) Small-bowel transit. (j) Distal colonic bead-expulsion time in control and SLICK-Hx*Csf1* mice.

To determine whether neuron-derived CSF1 contributes to MMs maintenance, we selectively deleted CSF1 from enteric neurons using SLICK-Hx*Csf1* mice following tamoxifen administration (**Figure 6b**). Immunohistochemical staining of the YFP reporter confirmed recombination within enteric neurons (**Supplementary Figure 7**), consistent with a recent report that SLICK-H drives recombination in enteric neurons but not enteric glia (10). Neuronal *Csf1* deletion significantly reduced MHCII⁺ MMs in the SI myenteric plexus, whereas colonic MMs abundance was unchanged (**Figures 6c,d**). Thus, enteric neurons-derived CSF1 provides regionally selective trophic support to MMs within the SI myenteric plexus.

We next asked whether disruption of neuron-derived CSF1 affected the enteric neuronal compartment. Neuronal *Csf1* deletion reduced both ANNA-1⁺ and nNOS⁺ neurons in the SI but not in the colon (**Figures 6c-f**). Consistent with the selective reduction of nNOS⁺ neurons and the preservation of ChAT⁺ neurons following systemic CSF1 neutralization (**Figure 5**; **Supplementary Figure 5**), these findings reveal a close association between the neuronal population enriched for *Csf1* expression and the neuronal population affected by disruption of the CSF1-dependent macrophage niche.

Finally, we examined the functional consequences of neuronal *Csf1* deletion. SLICK-Hx*Csf1* mice exhibited accelerated whole-gut transit (**Figure 6g**) and increased small-bowel transit (**Figure 6i**), whereas gastric emptying (**Figure 6h**) and distal colonic bead expulsion (**Figure 6j**) were unchanged. This regional physiological phenotype closely mirrored that observed following systemic CSF1 neutralization and occurred in the same intestinal region in which neuronal *Csf1* deletion reduced both MMs and nitrergic neurons.

Together, these findings identify a region- and neuronal subtype-specific trophic relationship between enteric neurons and MMs. In the SI, nNOS⁺ neurons are enriched for *Csf1* expression, and neuron-derived CSF1 provides trophic support to the CSF1-dependent MMs compartment. Loss of neuronal *Csf1* is accompanied by reductions in both MMs and nitrergic neurons and by altered small-intestinal motility. Combined with the selective effects of CSF1-dependent MMs depletion on nitrergic neurons (**Figure 5**), these findings support a reciprocal trophic relationship in which nitrergic neurons sustain the SI macrophage niche through CSF1, while CSF1-dependent MMs contribute to nitrergic neuronal homeostasis.

## DISCUSSION

Our study identifies regional specialization as a fundamental feature of intestinal MMs biology. MMs residing in different intestinal regions differed in their trophic requirements, responses to perturbation, recovery capacity, and functional relationships with the enteric nervous system. This heterogeneity became evident following transient CSF1 neutralization whereSI MMs were highly sensitive to CSF1 neutralization yet underwent robust repopulation, whereas colonic MMs were comparatively resistant to depletion but exhibited limited subsequent recovery. Moreover, restoration of macrophage abundance did not fully restore the steady-state macrophage state, as recovered populations remained transcriptionally remodeled and exhibited altered communication with the surrounding neuromuscular compartment. These observations broaden the concept of tissue-specific macrophage adaptation, revealing that substantial macrophage specialization can exist between anatomically and functionally distinct regions of the gut.

CSF1-CSF1R signaling is a major regulator of tissue macrophage survival (2), yet perturbation of this common trophic pathway produced markedly different outcomes across intestinal regions. A larger macrophage population persisted in the colon despite prolonged and high-dose CSF1 neutralization, but this relative resistance did not confer greater repopulation capacity. SI recovery was associated with persistence of macrophages labeled before depletion and preserved proliferative activity, whereas the colon showed reduced representation of the pre-depletion-labeled population, diminished proliferation, and persistent recruitment-associated remodeling. These divergent cellular trajectories suggest that regional MMs populations differ not only in their trophic requirements but also in the mechanisms contributing to their recovery. Importantly, robust numerical recovery in the SI did not fully restore steady-state transcriptional programs or predicted interactions with neurons, glia, ICCs, and stromal populations. Thus, macrophage abundance, cellular state, and functional integration can recover to different extents following transient perturbation.

The interaction between MMs and the enteric nervous system provides a functional example of this regional specialization. CSF1 neutralization selectively reduced SI nitrergic neurons without affecting cholinergic neurons and was accompanied by synapse remodeling and altered small-intestinal motility. Near-complete macrophage depletion through CSF1R blockade produced a distinct neuronal phenotype, with broader reductions in total and nNOS⁺ neuronal abundance but preservation of the relative representation of nitrergic neurons. Thus, the neuronal consequences of MMs perturbation did not simply scale with the magnitude of macrophage loss. Strikingly, *Csf1* expression was enriched in SI nNOS⁺ neurons but low or absent in ChAT⁺ neurons, and neuron-specific *Csf1* deletion reduced both SI MMs and nitrergic neurons while reproducing key features of the small-intestinal motor phenotype. Together, these findings identify a region- and neuronal subtype-specific relationship in which nitrergic neurons provide an important source of CSF1 for SI MMs, while CSF1-dependent MMs contribute to nitrergic neuronal homeostasis and intestinal function.

These findings challenge a model in which intestinal MMs are maintained through uniform trophic and neuroimmune mechanisms across the GI tract. Neuronal *Csf1* deletion did not significantly reduce colonic MMs, and a substantial colonic MMs population persisted during systemic CSF1 neutralization. This population remained sensitive to CSF1R blockade, suggesting that their maintenance may depend on alternative trophic signals acting through CSF1R. Together, these findings indicate that the contribution of individual trophic pathways to macrophage maintenance differs across intestinal regions. Identifying the trophic signals that regulate MMs in the colon and whether they are mediated by the enteric neurons or other cells in the ENS is an important task for future studies to enhance our understanding of the regional organization of MMs.

Several questions remain to be addressed, including the trophic signals, in addition to CSF1, supporting MMs maintenance in different intestinal regions, the mechanisms limiting colonic MMs recovery, and the contribution and long-term fate of recruited monocytes. The preferential localization of residual MMs near blood vessels following CSF1 neutralization raises the possibility that perivascular microenvironments contribute to MMs persistence, although the signals provided by the vascular environment and their functional significance remain unknown. The macrophage-derived signals contributing to nitrergic neuronal homeostasis also remain to be identified. Taken together, our findings show that intestinal MMs are regionally specialized in their trophic dependencies, recovery capacity, and functional interactions with the ENS. These findings also add to a growing body of evidence that macrophage identity is shaped not only by the organ of residence but also by anatomically and functionally distinct niches within a single organ, extending this principle, previously described in other organs, to the regional level of the gut and provide a framework for understanding how locally imprinted immune-neural circuits contribute to region-specific tissue homeostasis and region-specific vulnerability and repair in GI diseases.

## METHODS

### Mouse models

All procedures were carried out in accordance with the NIH Guide for the Care and Use of Laboratory Animals and approved by the Mayo Clinic Institutional Animal Care and Use Committee. B6.129P2(Cg)-*Cx3cr1^tm1Litt^*/J (CX_3_CR-1^GFP^), B6.Cg-*Gt(ROSA)26Sor^tm9(CAG-tdTomato)Hze^*/J (Ai9), STOCK Tg (Thy1-cre/ERT2,-EYFP) HGfng/PyngJ (SLICK-H^ER-Cre^), C57BL/6J (WT), B6.129P2(FVB)-*Lyz2^tm1(cre/ERT2)Grtn^*/J (*LysM*^CreERT2^), B6;129S6-*Chat^tm2(cre)low/^*J (ChAT^IRES-Cre^) and B6.Cg-Gt(ROSA) 26Sor^tm3(CAG-EYFP)Hze/^J (Ai3) strains were purchased from the Jackson Laboratory. *Csf1* flox mice were obtained from Dr. Jiang of UTHealth San Antonio, TX. To genetically label cholinergic neurons, ChAT^IRES-Cre^ were crossed with Ai3 mice to generate the ChATxAi3 mouse model. To genetically label myeloid cells, *LysM* ^CreERT2^ were crossed with Ai9 mice to generate the *LysM* ^CreERT2^xAi9 mouse model. To genetically delete CSF1 from neurons, SLICK-H^ER-Cre^ were crossed with *Csf1* flox mice to generate the SLICK-Hx*Csf1* mouse model. Mice were genotyped using the automated genotyping system Transnetyx (Cordova, TN, USA). Male mice, 6-8 weeks were used for the experiments. Male mice were used to minimize biological variability and maintain consistency across experimental groups. Water and food were available *ad libitum*, the lighting regime was a standard 12 h light and 12 h dark, temperature was maintained constant at 21 ± 2°C. Mice were humanely euthanized by carbon dioxide exposure followed by cervical dislocation in accordance with established institutional protocols.

### Tamoxifen-induced *Cre* recombination

Temporal control of cell-specific recombination was obtained by administering tamoxifen (10mg/ml in corn oil; Sigma-Aldrich, St. Louis, MO) to 6-week-old mice via daily intraperitoneal doses (75mg/kg) for three consecutive days. All experiments were performed after a 1-week tamoxifen washout. Vehicle-treated (corn oil) *LysM* ^CreERT2^xAi9 mice were used to validate tamoxifen-dependent reporter activation, whereas tamoxifen-treated mice not subjected to αCSF1 depletion served as controls for lineage-labeling recovery experiments.

### Muscularis Macrophage depletion

To deplete muscularis macrophages, wild-type (WT) mice (6-8 weeks) were treated with either a CSF1-neutralizing antibody (8.5 or 17 µg/g/day; Bio X Cell, InVivoMAb anti-mouse CSF1, #BE0204) administered daily for 7 days or a CSF1R neutralizing antibody (37.5 µg/g; Bio X Cell, InVivoMAb anti-mouse CSF1R, #BE0213) administered twice over a 7-day period (4). Control mice for both neutralizing antibody experiments received intraperitoneal (IP) injections of an isotype control antibody (Bio X Cell, InVivoMAb rat IgG1 anti-horseradish peroxidase [HRP], BE0088) matching the dose and administration schedule of the corresponding treatment group.

### In vitro primary culture of muscularis macrophages

Mucosal-free muscularis externa from small intestine and colon of Cx3Cr1^GFP+^ mice were dissociated as previously described (11). Cell sorting was performed using a BD FACSMelody™ Cell Sorter with BD FACSChorus™ Software (BD Bioscience), located at the Mayo Clinic Flow Cytometry Core Facility (Rochester, MN). FACS-sorted CX3CR1^GFP⁺^ cells were seeded in 24-well plates at a density of 50,000 cells/well and cultured in complete media (DMEM/F-12 supplemented with 10% FBS, 1% antibiotic-antimycotic, and 20 ng/mL mouse CSF1 (ProSpec, #CYT-439)) for 3 days. After an attachment period and for the rest of the experiment 50% of media was replaced daily. Cells were left to grow and differentiate for a total of 7 days after the attachment period. Media was completely replaced with αCSF1- or IgG-enriched complete media after differentiation period and sustained during a 7-day treatment period. When indicated, cells were allowed to recover with antibody-free complete media for at least 10 days. After the attaching period cells were followed daily by live-cell confocal fluorescence microscopy to assess morphology and cell counts. Each data point represents the average of 3 randomly selected regions of the same well.

### Single-cell sequencing

Single-cell sequencing was performed on the muscularis externa obtained from the small intestine and colon of age-matched mice from the different experimental groups. Cells were assessed for viability using the Vi-Cell XR Cell Viability Analyzer (Beckman-Coulter), and their count was determined. Following this, barcoded Gel Beads were thawed, and a cDNA master mix was prepared according to Chromium Single Cell 3’ v3 library kit instructions. Cells were mixed with the master mix and Gel Beads, then processed using the Chromium Controller to capture individual cells in uniquely labeled Gel Beads-In-Emulsion (GEMs). After collecting the GEMs, reverse transcription, dissolution, and cDNA clean-up were performed, resulting in a pool of barcoded cDNA. This cDNA was then used to construct libraries, incorporating Illumina sequencing primers and unique i7 Sample index, and assessed for quality using Qubit High Sensitivity assays (Thermo Fisher Scientific), Agilent Bioanalyzer High Sensitivity chips (Agilent), and Kapa DNA Quantification reagents (Kapa Biosystems). Sequencing was conducted at a rate of 60,000 fragment reads per cell according to Illumina’s standard protocol, using the Illumina cBot and HiSeq 3000/4000 PE Cluster Kit. The flow cells were sequenced as 100 X 2 paired end reads on an Illumina HiSeq 4000 using HiSeq 3000/4000 sequencing kit and HCS v3.3.52 collection software. Base-calling was performed using Illumina’s RTA version 2.7.3. Raw BCL files were demultiplexed into FASTQ sequencing files using the Cell Ranger version 3.1 suite from 10xGenomics.

### Single cell RNA-seq preprocessing and quality control

We used 10X Genomics Cell Ranger Single Cell Software Suite (v7.2.0) to perform alignment to mm10 reference genome, filtering, barcode counting, and UMI counting. For subsequent analysis, we followed the standard integrated analysis workflow in the Seurat package (v4)(12). Cell doublet identification was performed using scDblFinder (13). Predicted doublets were removed from downstream analysis. The remaining cells that met any of the criteria below were further removed from downstream analysis for each sample: 1). Cells expressing fewer than 50 genes; 2) Cells with a total UMI lower than 200; 3). Cells with percentage of mitochondrial reads > 40%; 4). Cells with a percentage of hemoglobin concentration higher than 20%. Genes expressed in fewer than 10 cells were removed from downstream analysis. Each sample was normalized using SCTransform and scaled for each gene across all cells (14). Control samples from the small intestine and colon were first integrated and clustered on the low-dimensional space with resolution 0.3 to identify 23 unsupervised clusters. Samples were then mapped to the integrated object using Seurat’s FindTransferAnchors.

### Single-cell RNA-sequencing analyses

#### Data processing and preparation

Raw sequencing data were processed using the 10x Genomics Cell Ranger Single Cell Software Suite (v7.2.0) (15), including alignment to the mm10 mouse reference genome, cell-barcode filtering and UMI quantification. Subsequent analyses and quality control were performed using Seurat (v5) (16). Cell doublets were identified using scDblFinder and excluded from downstream analyses. Cells were additionally excluded if they expressed fewer than 50 genes, contained fewer than 200 total UMIs, had >40% mitochondrial transcripts, or >20% hemoglobin-associated transcripts. Genes detected in fewer than 10 cells were also excluded. Following quality control, gene-expression data from each sample were normalized using SCTransform to reduce technical variation, including differences in sequencing depth between cells.

#### Cell-lineage annotation and macrophage subtyping

Samples were integrated and subjected to unsupervised clustering at a resolution of 0.8 to define reference cell populations. Broad cellular populations were annotated using canonical lineage markers and cell-type scores. Marker sets included endothelial cells (*Pecam1, Vwf, Kdr, Cdh5*), epithelial cells (*Epcam, Krt8, Krt18, Krt19*), erythrocytes (*Hbb-bs, Hbb-bt, Hba-a1, Hba-a2, Alas2*), glial cells (*S100b, Plp1, Sox10*), interstitial cells of Cajal (ICCs; *Kit, Ano1*), macrophages (*Lyz2, Tyrobp, Csf1r, Adgre1, C1qa*), mesothelial cells (*Msln, Wt1, Upk3b*), fibroblasts (*Col1a1, Col1a2, Dcn, Lum*), myofibroblasts (*Acta2, Tagln, Col1a1*), neurons (*Tubb3, Snap25, Syt1*), pericytes (*Rgs5, Pdgfrb, Cspg4*), smooth-muscle cells (*Acta2, Tagln, Myh11, Cnn1*), T cells (*Cd3d, Cd3e, Trac*) and vascular smooth-muscle cells (*Myh11, Notch3, Tagln*). Cell-type scores derived from these marker sets were used to support cell-lineage assignment.

Macrophage-containing clusters were isolated and re-clustered to resolve their heterogeneity. Dimensionality reduction was performed using principal component analysis (PCA). Thirty principal components were calculated, of which the first 20 were used to construct a shared nearest-neighbor graph and perform graph-based clustering at a resolution of 0.8. The same 20 principal components were used to generate macrophage-specific UMAP representations.

To examine changes in MM population composition across experimental conditions, macrophages were assigned to three major populations based on canonical marker expression (9). T cell immunoglobulin and mucin domain containing 4, lymphatic vessel endothelial hyaluronan receptor 1, folate receptor beta (TLF) macrophages were identified by detectable expression of at least one of *Timd4, Lyve1 or Folr2*. Among cells not classified as TLF, C-C motif chemokine receptor 2 (CCR2) macrophages were identified by detectable *Ccr2* expression. Among macrophages classified as neither TLF nor CCR2, an MHCII expression score was calculated as the mean normalized expression of *H2-Eb1, H2-Aa, H2-Ab1* and *Cd52*. Cells with scores at or above the 20th percentile of the eligible-cell distribution were classified as MHCII macrophages.

#### UMAP visualization and macrophage composition

Cellular identities were visualized using Uniform Manifold Approximation and Projection (UMAP) projections. Whole-tissue UMAPs were used to display the distribution of major cellular lineages, whereas macrophage-specific UMAPs were used to visualize TLF, CCR2 and MHCII macrophages. UMAPs were additionally stratified by experimental conditions to compare cellular distributions across control, treated and recovery samples. Expression of selected genes and transcriptional program scores was overlaid onto the UMAP representations.

Changes in macrophage composition were assessed separately in the colon and small intestine. For each experimental condition, the abundance of TLF, CCR2 and MHCII macrophages was expressed as a proportion of the total macrophage population within the corresponding tissue. Differences in subtype proportions across conditions were evaluated using proportion tests, followed by pairwise comparisons where appropriate. Subtype proportions were also compared between colon and small intestine within each experimental condition. Pairwise P values were adjusted for multiple testing using the Benjamini-Hochberg procedure to control the false discovery rate (FDR).

#### Differential gene-expression analysis and cross-tissue comparison

Differential gene-expression analysis was performed separately for each macrophage subtype and tissue using Seurat v5. Pairwise comparisons were performed for control versus treated, control versus recovery and treated versus recovery using the Wilcoxon rank-sum test.

Genes expressed in at least 10% of cells in either comparison group were tested. Comparisons were performed only when at least three cells were available in each group. Genes were considered differentially expressed when the adjusted P value was <0.05 and the absolute average log2 fold change was ≥0.25. Volcano plots were used to display the magnitude and statistical significance of transcriptional changes.

To determine whether transcriptional responses were shared between intestinal regions, differentially expressed genes from colon and small intestine were compared for the same macrophage subtype and experimental condition. Only genes measurable in both datasets were included. The degree of overlapping was quantified using the Jaccard index, and enrichment of overlapping genes was evaluated using Fisher’s exact test followed by FDR correction. Upregulated and downregulated genes were additionally considered separately. For genes significant in both tissues, directional concordance was quantified as the proportion of genes showing the same direction of effect and by Spearman correlation between colon and small-intestinal log2 fold-change estimates.

To investigate potential cellular sources of macrophage-supporting signals, expression of the CSF1R ligand *Csf1* was examined across major cellular lineages in the complete colon and small-intestinal datasets. For each lineage and experimental condition, the proportion of cells expressing each gene and the mean normalized expression level were calculated. Expression patterns were visualized using lineage-stratified summaries and UMAP projections to assess tissue- and condition-specific differences in the cellular sources of *Csf1*. *Csf1* expression was specifically evaluated in nNOS⁺ and ChAT⁺ neuronal populations, classified according to expression of *Nos1* and *Chat*, respectively.

#### Functional transcription programs and heatmap analysis

Macrophage transcriptional states were characterized using six predefined functional gene programs: (i) homeostatic identity (*Tyrobp, Cx3cr1, Csf1r, Mertk, Cd163*); (ii) resident/homeostatic function (*Apoe, Psap, Ms4a4a, Fcrls, Stab1*); (iii) neuro-support and trophic signaling (*Igf1, Bmp2, Psap, Grn, Apoe*); (iv) antigen presentation (*H2-Aa, H2-Ab1, H2-Eb1, Cd74, Cd86, Itgam*); (v) inflammatory/NF-κB activation (*Nfkbia, Rel, Junb, Tnf, Ccl2, Cxcl10, Mmp9*); and (vi) inflammasome/cytokine signaling (*Nlrp3, Casp4, Il1b, Tnf, Il18*). Only genes detected in the corresponding dataset were included.

Functional-program scores were calculated from the expression of the genes comprising each signature and evaluated separately for each macrophage subtype within the colon and small intestine. Treated and recovery cells were each compared with control cells using Wilcoxon rank-sum tests, with P values adjusted for FDR. Changes in each program were summarized as the difference in mean score between conditions (Δ mean score).

Gene-level expression patterns within the functional programs were additionally visualized using heatmaps. For each gene, normalized expression was first averaged across all cells within each tissue and experimental condition. These mean expression values were then standardized separately for each gene across the six tissue-by-condition groups-colon-control, colon-treated, colon-recovery, small intestine-control, small intestine-treated and small intestine-recovery-to generate Z-scores. Positive values therefore indicated expression above that gene’s overall mean across groups, whereas negative values indicated expression below it. Colon and small-intestinal heatmaps were displayed separately using the same gene-wise standardization and color scale.

#### Proliferation and recruitment programs

Macrophage proliferative and recruitment-associated states were quantified using two additional predefined transcriptional signatures. The proliferation signature comprised *Mki67*, *Top2a*, *Stmn1*, *Hmgb2*, *Pcna*, *Cdk1*, *Ube2c* and *Birc5*. The recruitment/inflammatory-monocyte signature comprised *Ccr2*, *Ly6c2*, *Plac8*, *S100a8*, *S100a9*, *Il1b*, *Fn1*, *Tgfbi*, *Vcan* and *Ccl2*. Only genes detected in the corresponding dataset were included. Single-cell proliferation and recruitment scores were calculated from the genes comprising each signature and evaluated separately in colon and small intestine across TLF, CCR2 and MHCII macrophages. Differences among groups were evaluated using Kruskal-Wallis tests followed, where appropriate, by pairwise Wilcoxon rank-sum tests, with P values adjusted for FDR. Individual signature genes and transcriptional scores were visualized across macrophage populations and experimental conditions using grouped expression plots and UMAP projections.

#### Cell-cell communication analysis

Cell-cell communication was inferred using CellChat (v2.2.0.9001) (17, 18) and the mouse ligand-receptor interaction database. Colon and small intestine were analyzed separately within each experimental condition, generating independent communication networks for control, treated and recovery samples.

Macrophages were considered as a single population, while the other major cellular lineages were retained as potential signaling partners. Cell populations represented by fewer than 10 cells within a given tissue and condition were excluded from the analysis. Communication networks were inferred from normalized gene-expression data. Overexpressed signaling genes and ligand-receptor interactions were identified, and expression values were smoothed using the mouse protein-protein interaction network before estimating communication probabilities. Communication probabilities were estimated independently of cell-population size, so that inferred signaling strength reflected ligand-receptor expression rather than differences in cell abundance. Pathway-level and aggregate communication networks were subsequently derived from the inferred ligand-receptor interactions.

Macrophage-centered interactions were classified as outgoing when macrophages acted as the signaling source and incoming when macrophages acted as the target. Macrophage self-interactions were excluded from partner-specific analyses. Communication with fibroblasts, glial cells, interstitial cells of Cajal, pericytes and neurons was examined in detail.

For each ligand-receptor pair, communication strength was summarized as the summed inferred communication probability (sum_prob) for each tissue, condition, communication direction and partner cell type. To identify the most dynamically changing signals, ligand-receptor pairs were ranked according to the absolute change in summed communication probability between recovery and control within each macrophage-partner comparison. The highest-ranking interactions were subsequently followed across control, treated and recovery conditions.

#### Whole-mount Immunohistochemistry (IHC)

Stomachs, small intestines and colons were collected, and the muscularis externa separated from the mucosa. Tissues were then fixed with 4% paraformaldehyde (PFA, Sigma Aldrich) in 0.1 M phosphate buffer overnight at 4°C. After fixation, following three washes to remove residual PFA, tissues were incubated with a blocking solution containing 5% Bovine Serum Albumin (BSA, Sigma Aldrich) in 1x Phosphate-buffered saline (PBS, Sigma Aldrich) and 1% TritonX-100 (Sigma Aldrich) overnight at room temperature (RT). After blocking, tissues were labeled in the presence of the primary antibodies overnight at RT (**Table 1**). After washing, the tissues were incubated with the appropriate secondary antibodies (**Table 2**) overnight at RT, in the dark, and then mounted using SlowFade™ Gold Antifade Mountant (Thermo Fisher Scientific). For all immunohistochemical studies, controls omitting the primary antibodies and controls in double labeling experiments with a mis-matched secondary antibody were performed. For quantitative analysis, labeled tissues were examined with a laser scanning confocal microscope (Nikon AX-R) using a 20X (NA 0.75 WI) Nikon Apo LWD objective (Nikon) using identical fluorescence acquisition settings (laser power, gain, and exposure) across all groups. Stacks of confocal images across the full thickness of the muscularis propria were obtained from 3-5 non-adjacent fields of view selected randomly per tissue/mouse. All the confocal image stacks were flattened into projections using NIS elements AR (Nikon) for quantification of the labeling, ensuring that signal from the entire z-depth was included in the analysis. Cell numbers were determined by manual counting. The number of macrophages is reported as percentage relative to the average number of cells in control condition.

**Table 1:** Primary antibodies.

| Name | Dilution | Experiment | Manufacturer | Cat. No. | Fluorochrome | RRID# |
| --- | --- | --- | --- | --- | --- | --- |
| ANNA-1 | 1:5000 | IF | - | Gift from Dr.<br>V. Lennon,<br>Mayo Clinic,<br>Rochester,<br>MN | - | - |
| MHC Class II<br>(MHCII) | 1:1000 | IF | eBioscience | 14-5321-81 | - | AB_467560 |
| Neuronal Nitric<br>Oxide Synthase<br>(NOS1) | 1:1000 | IF | Millipore | AB5380 | - | AB_91824 |
| Ki67 | 1:250 | IF | Abcam | ab16667 | - | AB_302459 |
| Yellow fluorescent<br>protein (YFP) | 1:500 | IF | Aves Lab | GFP-1010 | - | AB_2307313 |
| PECAM-1 | 1:250 | IF | R&D | AF3628 | - | AB_2161028 |
| Synaptophysin<br>(SYP) | 1:500 | IF | Abcam | ab16659 | - | AB_443419 |
| Anti-mouse<br>CD16/32 | 1:100 | FACS | Biolegend | 101301 | NA | AB_2794188 |
| Anti-mouse CD45<br>(30-F11) | 1:200 | FACS | Biolegend | 103183 | Spark Blue 574 | AB_2904274 |
| Anti-mouse CD11b<br>(M1/70) | 1:160 | FACS | ThermoFisher | 48-0112-82 | eFlour 450 | AB_1582236 |
| Anti-mouse CD115<br>(CSF1R) | 1:333 | FACS | ThermoFisher | 46-1152-80 | PerCP-eFluor 710 | AB_1059774<br>1 |

IF, Immunofluorescence; FACS, fluorescence-activated cell sorting.

**Table 2:** Secondary antibodies.

| **Secondary antibody** | **Supplier** | **Host** | **Clonality** | **Cat.No.** | **RRID#** |
| --- | --- | --- | --- | --- | --- |
| Cy3 anti-goat | Jackson<br>ImmunoResearch | Donkey | Polyclonal | 705-165-47 | - |
| Alexa Fluor 488 anti-rat | Jackson<br>ImmunoResearch | Donkey | Polyclonal | 712-545-150 | AB_2340683 |
| Cy5 anti-rat | Jackson<br>ImmunoResearch | Donkey | Polyclonal | 712-175-150 | AB_2340671 |
| Cy3 anti-rabbit | Jackson<br>ImmunoResearch | Donkey | Polyclonal | 711-165-152 | AB_2307443 |
| Cy5 anti-human | Jackson<br>ImmunoResearch | Donkey | Polyclonal | 709-175-149 | AB_2340539 |
| Cy3 anti-human | Jackson<br>ImmunoResearch | Goat | Polyclonal | 109-165-170 | AB_2810895 |

#### RT-qPCR

Total RNA was extracted from the muscularis externa of the stomach, small intestine, and colon using the RNeasy plus universal kit (Qiagen) according to the manufacturer’s specifications. RNA concentration was determined by using a NanoDrop ND-1000 spectrophotometer (Thermo Fisher Scientific). For first-strand cDNA synthesis, 500 ng of total RNA from each sample was reverse-transcribed using the SuperScript VILO complementary DNA Synthesis Kit (Invitrogen). Real-time PCR was performed using the LightCycler® 480 SYBR Green I Master Mix (Roche Molecular Systems). Primers were designed based on the mouse GenBank sequences (**Table 3**). The relative expression of mRNA was normalized by glyceraldehyde-3-phosphate dehydrogenase (Gapdh, Qiagen, catalog # PPM02946E-200) and calculated by the 2^−ΔΔCt^ method.

**Table 3:**
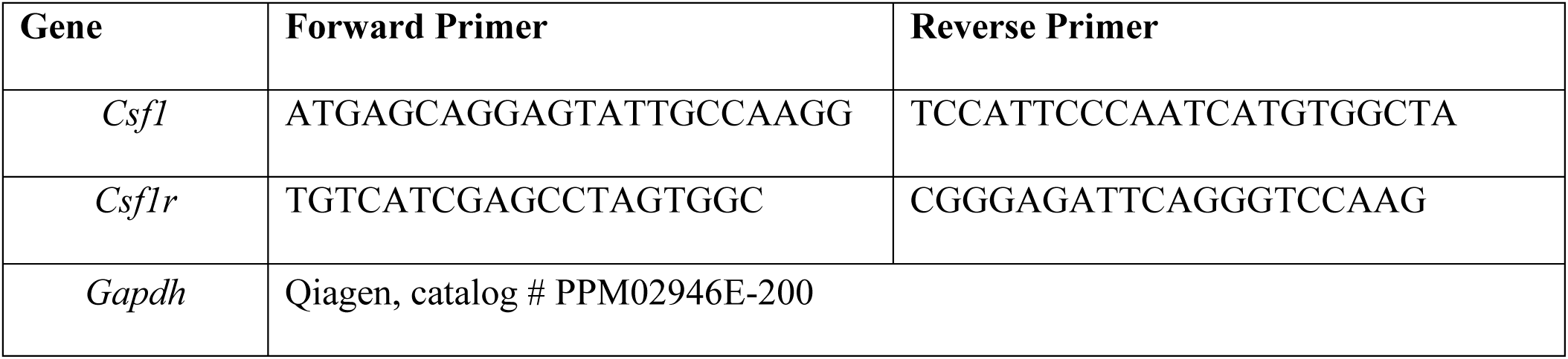
Primer sequences.

#### Flow cytometry

For lineage-labeling experiments, single-cell suspensions were prepared from the peripheral blood of *LysM*^CreERT2^xAi9 mice after single dose of Tamoxifen injection. Following exclusion of debris and doublets, tdTomato fluorescence was assessed to confirm tamoxifen-dependent labeling. Vehicle-treated *LysM*^CreERT2^xAi9 mice were used to establish background fluorescence and the tdTomato-positive gate. The antibodies used are listed in **Table 1**.

#### Whole gut transit (WGT)

Whole-gut transit was measured as previously described [17]. Briefly, mice received Carmine Red solution (6% w/v; Sigma-Aldrich) prepared in methylcellulose (0.5% w/v; Sigma-Aldrich) by oral gavage. Following gavage, mice were individually housed in a controlled environment and continuously monitored by video recording. Food and water were withheld for the duration of the experiment, which lasted up to approximately 8 h. Whole-gut transit time was defined as the interval between Carmine Red administration and the appearance of the first red fecal pellet.

#### Gastric emptying

Gastric emptying breath test was performed as described previously (19). Briefly, mice were fasted overnight with free access to water in a metabolic cage. Mice were then placed in a chamber flushed with CO2-free air. After a baseline reading, mice were fed 200 mg scrambled cooked egg yolk containing 2.5 μmol [13C] octanoic acid (Cambridge Isotope Laboratories, Tewksbury, MA). Finally, air containing exhaled breath was collected and analyzed to determine the 13C-to-12C ratio, represented as T half value. Each mouse was tested once a week for 4 consecutive weeks. The average T half value from 3 tests was used to calculate group mean values.

#### Small Bowel Transit

Small Bowel transit was measured as previously described (20). Briefly, mice were fasted for 4 hours providing access to water. 200 µL of fluorescent dye (rhodamine isothiocyanate (RITC)-dextran) (Sigma Aldrich) were administered by gavage. Mice were euthanized after 30 minutes using carbon dioxide inhalation, followed by cervical dislocation to ensure successful euthanasia. The GI tract was removed from mice and imaged with an in vivo imaging system (IVIS) to analyze the fluorescence distribution. Geometric center of the distribution was calculated using image J and reported as distance traveled by the dye.

#### Colonic transit

Colonic transit was assessed using the bead expulsion assay as previously described (21). Briefly, mice were briefly anesthetized with isoflurane, and a 3-mm glass bead was gently inserted 2 cm into the distal colon using a lubricated applicator. Following bead insertion, each mouse was individually placed in a clean cage, and the latency to bead expulsion was recorded. Mice that did not expel the bead within 100 minutes were assigned the maximum experimental time of 100 minutes. Colonic transit was expressed as the time, in minutes, required for bead expulsion.

### Statistical analysis

Data points represent biological replicates. Results are presented as means ± SEM. Statistical analyses were conducted using the GraphPad Prism 10.6.1 software (GraphPad, Boston, MA). For comparisons among multiple groups, one-way analysis of variance (ANOVA) followed by Bonferroni’s multiple comparison test was performed. Unpaired 2-tailed Student’s *t*-test was used for comparisons between two groups. ∗, ∗∗, ∗∗∗, and ∗∗∗∗ indicate statistical significance with *P* < 0.05, *P* < 0.01, *P* < 0.001, and *P* < 0.0001, respectively. Statistically nonsignificant results were labeled as ns.

**Supplementary Figure 1.**
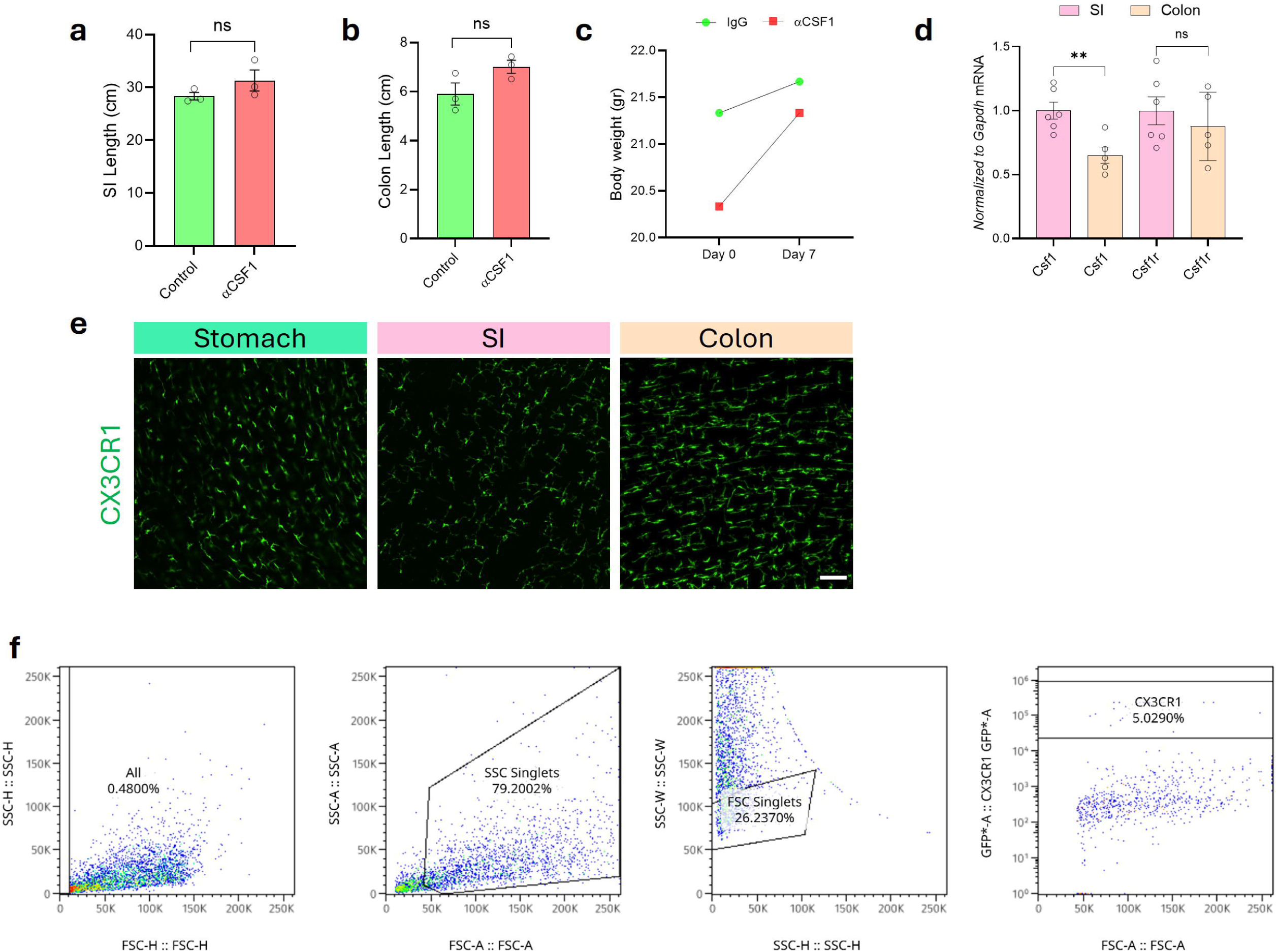
Validation of regional macrophage responses to CSF1 neutralization. (a) Small-intestinal length in control and αCSF1-treated mice. (b) Colonic length in control and αCSF1-treated mice. (c) Body weight before and after 7 days of control or αCSF1 treatment (*n =* 3 per group). (d) Relative *Csf1* and *Csf1r* expression in SI and colonic muscularis tissue measured by RT-qPCR. (e) Representative whole-mount images of CX3CR1-GFP⁺ MMs in the stomach, SI, and colon. (f) Representative flow-cytometry gating strategy used for the isolation of CX3CR1-GFP⁺ MMs from the intestinal muscularis. Bar graphs represent the mean ± SEM; ∗∗ *P* < 0.01, ns, nonsignificant (unpaired *t*-test). Scale bar: 100 μm.

**Supplementary Figure 2.**
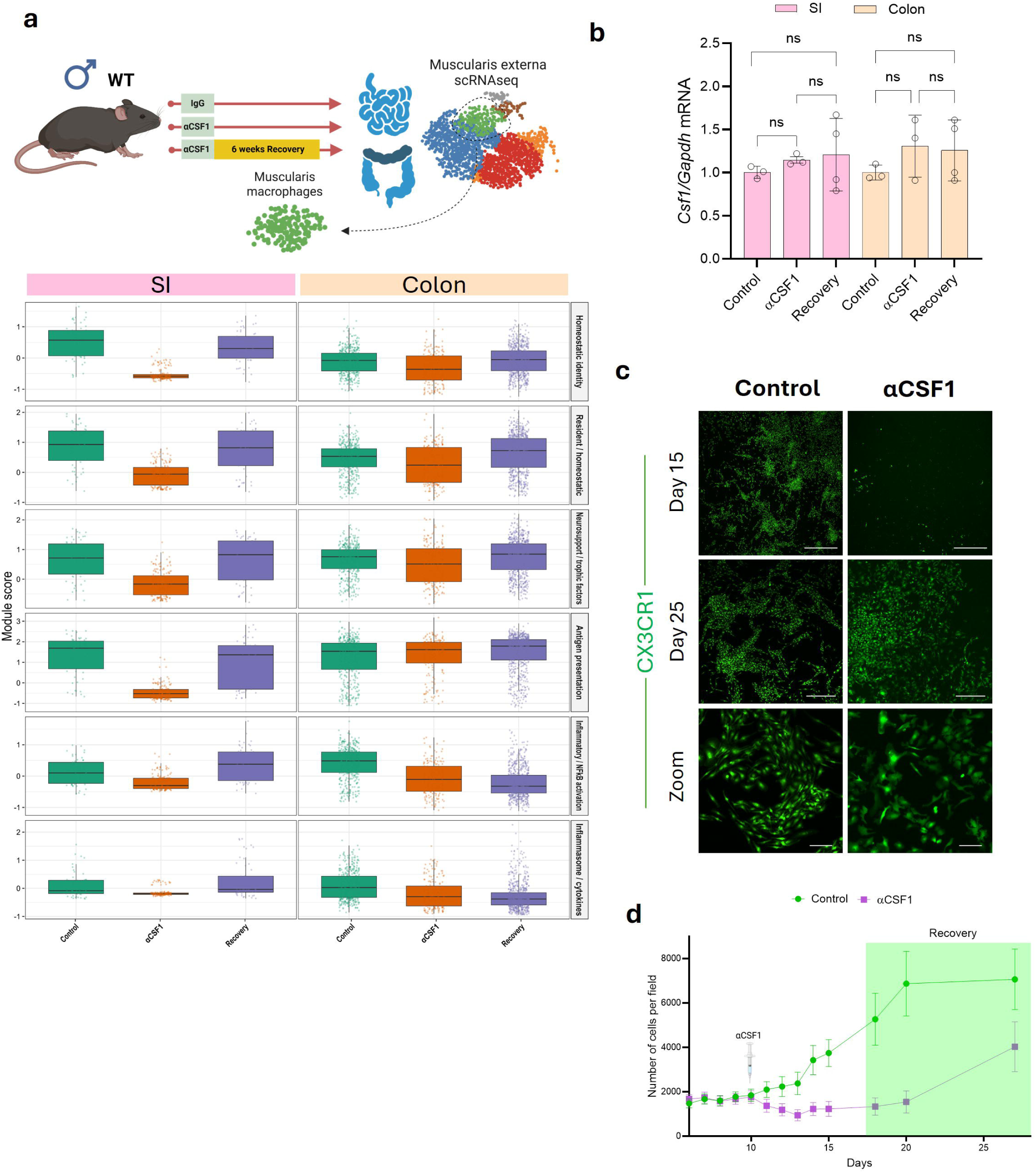
Macrophages surviving transient CSF1 neutralization retain the capacity to expand following antibody withdrawal. (a) Extended analysis of macrophage transcriptional programs across control, αCSF1-treated, and recovery conditions in the SI and colon. (b) Relative *Csf1* expression in SI and colonic muscularis externa across control, αCSF1-treated, and recovery conditions. (c) Representative images of CX3CR1-GFP⁺ MMs during ex vivo control, αCSF1 treatment, and recovery following withdrawal of αCSF1. (d) Quantification of GFP⁺ macrophage abundance during αCSF1 treatment and following antibody withdrawal. Each point represents one biological replicate unless otherwise indicated. Bar graphs represent the mean ± SEM; ns, nonsignificant (one-way ANOVA with Bonferroni’s multiple comparison). Scale bar: 500 μm.

**Supplementary Figure 3.**
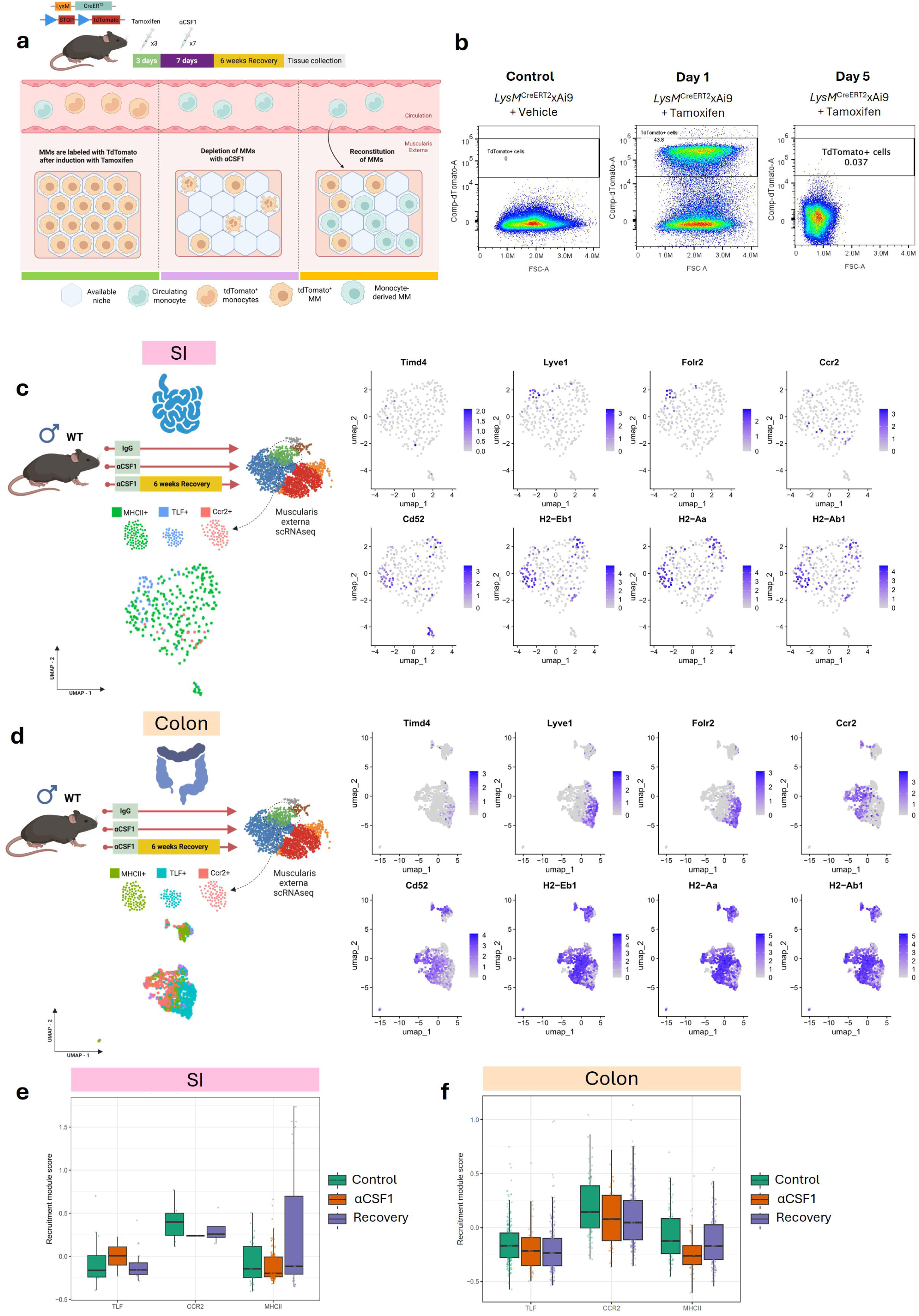
Validation and transcriptional characterization of macrophage populations during recovery. (a) Schematic representation of the inducible lineage-labeling strategy used to label macrophages before αCSF1-mediated depletion and assess their contribution to the macrophage compartment following recovery. (b) Representative flow-cytometry plots showing tdTomato labeling of peripheral blood cells from vehicle- and tamoxifen-treated *LysM* ^CreERT2^xAi9 mice. (c) UMAP visualization of SI macrophage populations and expression of *Timd4*, *Lyve1*, *Folr2*, *Ccr2*, *Cd52*, *H2-Eb1*, *H2-Aa*, and *H2-Ab1* used to cluster macrophages into MHCII^+^, TLF^+^, and CCR2^+^ subpopulations. (d) UMAP visualization of colonic macrophage populations and expression of the corresponding macrophage markers. (e) Recruitment module score across SI macrophage populations under control, αCSF1-treated, and recovery conditions. (f) Recruitment module score across colonic macrophage populations under control, αCSF1-treated, and recovery conditions.

**Supplementary Figure 4.**
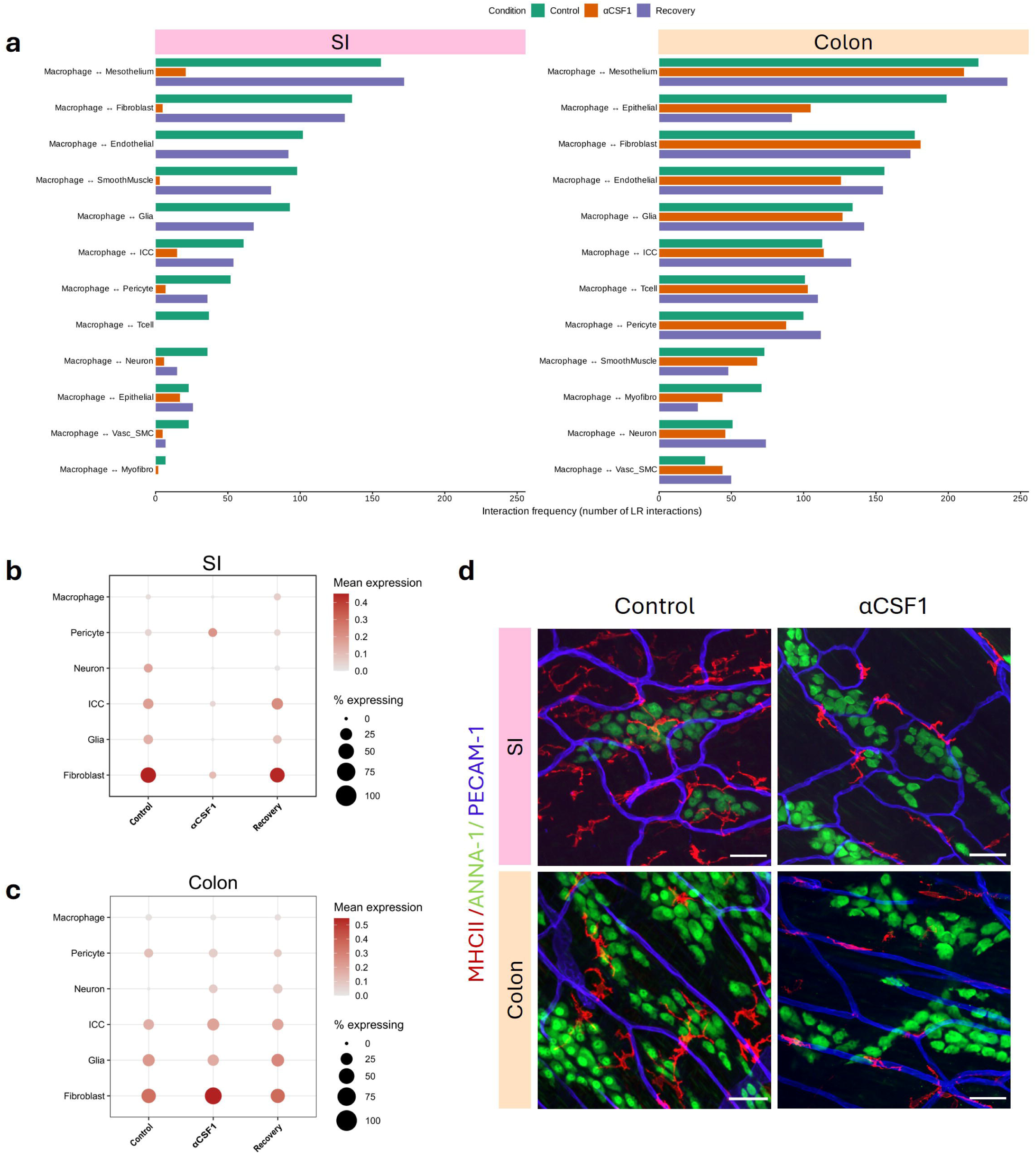
Regional remodeling of macrophage communication and spatial organization following CSF1 neutralization. (a) Extended analysis of inferred macrophage interactions with muscularis externa cell populations across control, αCSF1-treated, and recovery conditions in the colon and SI. Bars indicate the interaction frequency between MMs and the indicated cell populations. (b) *Csf1* expression across major SI cellular populations under control, αCSF1-treated, and recovery conditions. Dot size indicates the percentage of cells expressing *Csf1*, and color indicates mean expression. (c) *Csf1* expression across major colonic cellular populations under control, αCSF1-treated, and recovery conditions. (d) Representative whole-mount images of MHCII⁺ MMs, ANNA-1⁺ enteric neurons, and PECAM1⁺ blood vessels in the SI and colon from control and αCSF1-treated mice.

**Supplementary Figure 5.**
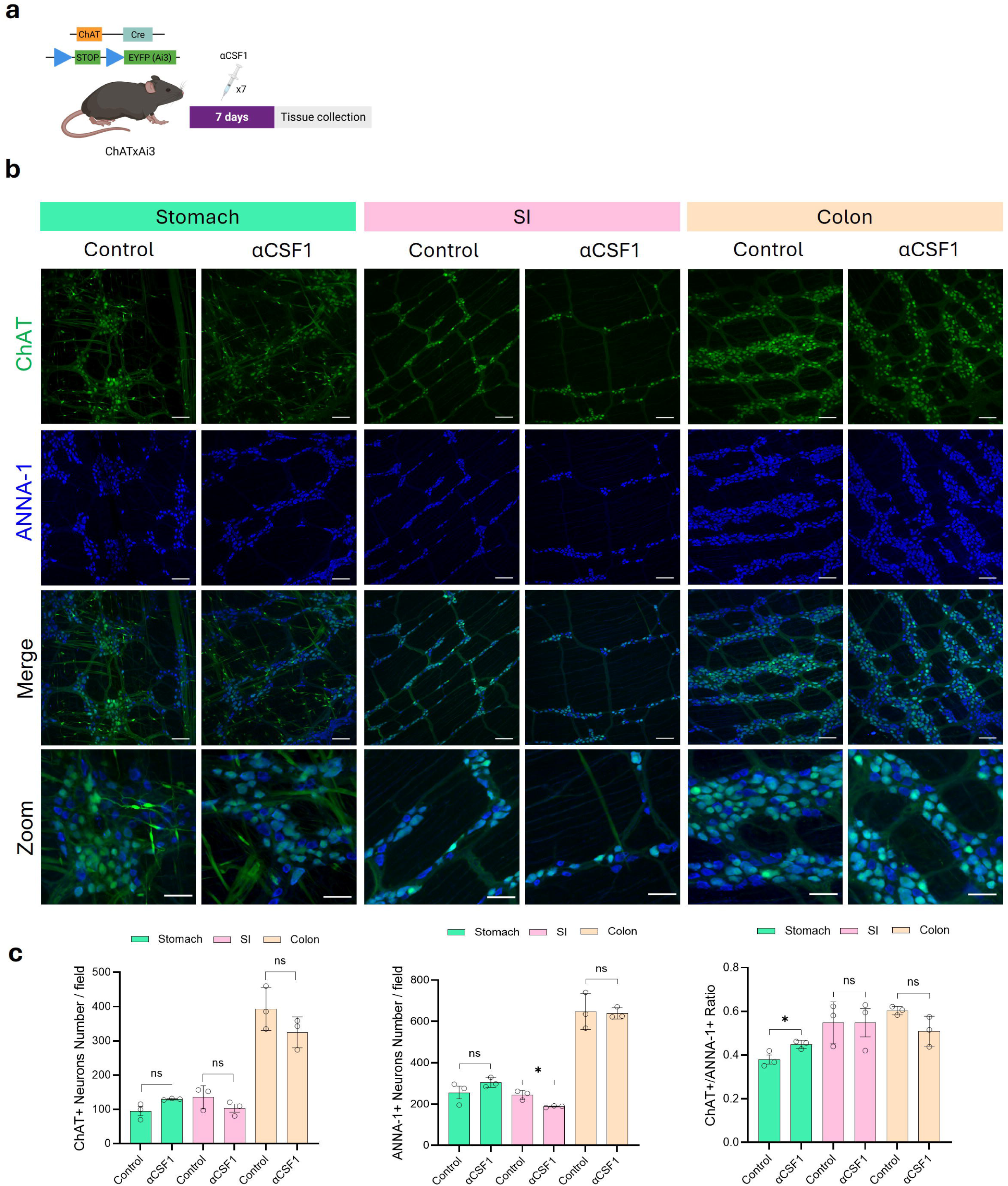
Cholinergic enteric neurons are preserved following CSF1 neutralization. (a) Schematic representation of αCSF1 treatment in ChATxAi3 reporter mice. (b) Representative whole-mount images of ChAT^+^ and ANNA-1⁺ enteric neurons in the stomach, SI, and colon from control and αCSF1-treated mice. (c) Quantification of ChAT⁺ neurons, total ANNA-1⁺ neurons, and the ChAT⁺/ANNA-1⁺ neuronal ratio in the stomach, SI, and colon following αCSF1 treatment.

**Supplementary Figure 6.**
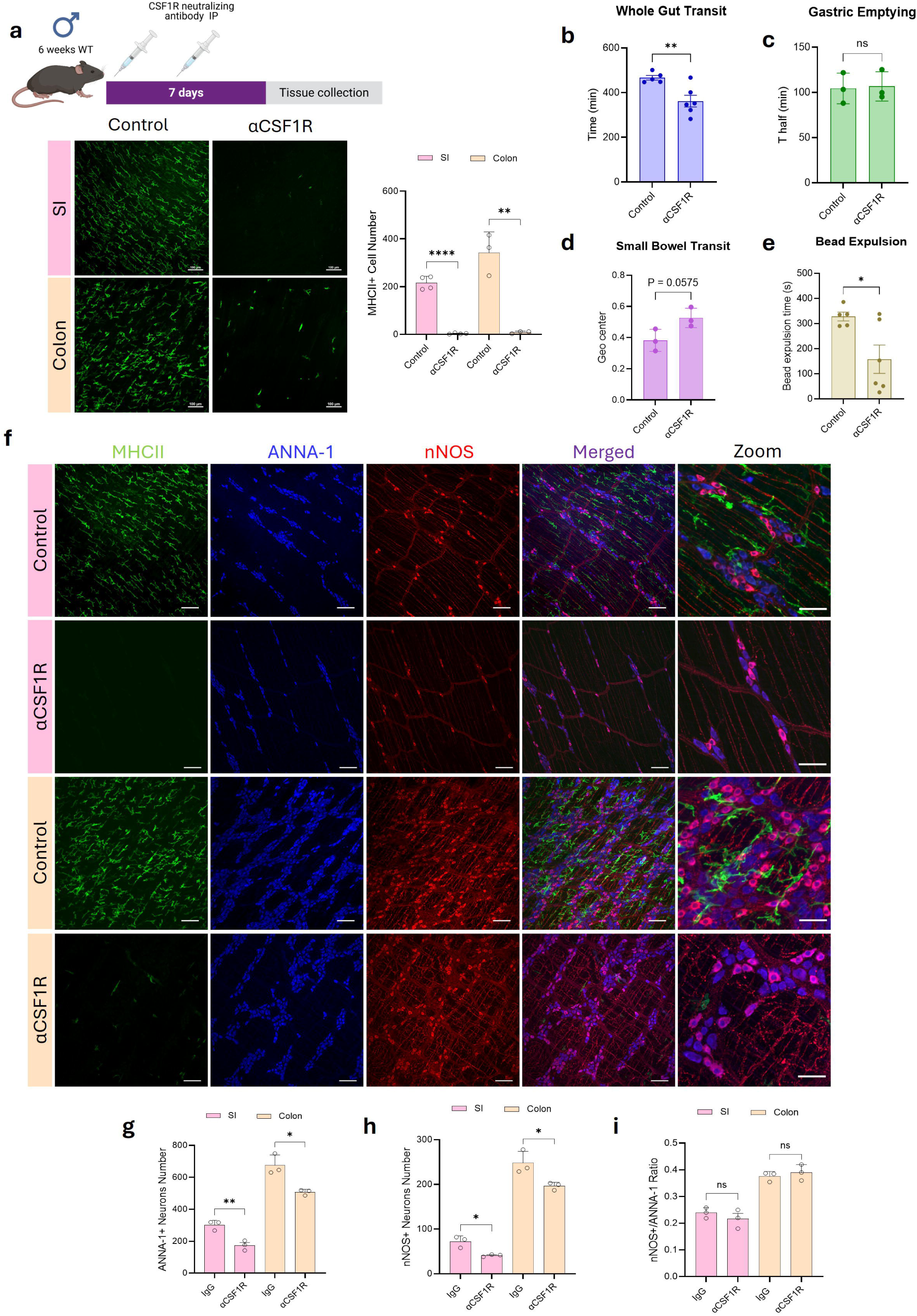
CSF1R neutralization produces near-complete MM depletion and alters gastrointestinal motility. (a) Schematic representation of αCSF1R treatment, with representative whole-mount images and quantification of MHCII⁺ MMs in the SI and colon from control and αCSF1R-treated mice. (b) Whole-gut transit time. (c) Gastric emptying. (d) Small-bowel transit. (e) Distal colonic bead-expulsion time in control and αCSF1R-treated mice. (f) Representative whole-mount images of MHCII⁺ MMs, ANNA-1⁺ enteric neurons, and nNOS⁺ neurons in the SI and colon from control and αCSF1R-treated mice. (g) Quantification of total ANNA-1⁺ enteric neurons. (h) Quantification of nNOS⁺ enteric neurons. (i) Proportion of nNOS⁺ neurons to total ANNA-1⁺ neurons in the SI and colon following αCSF1R treatment. Each point represents one biological replicate unless otherwise indicated. Bar graphs represent the mean ± SEM; ∗ *P* < 0.05, ∗∗ *P* < 0.01, ∗∗∗* *P* < 0.0001, ns, nonsignificant (unpaired *t*-test). Scale bar: 100 μm (a,f), 50 μm (zoom).

**Supplementary Figure 7.**
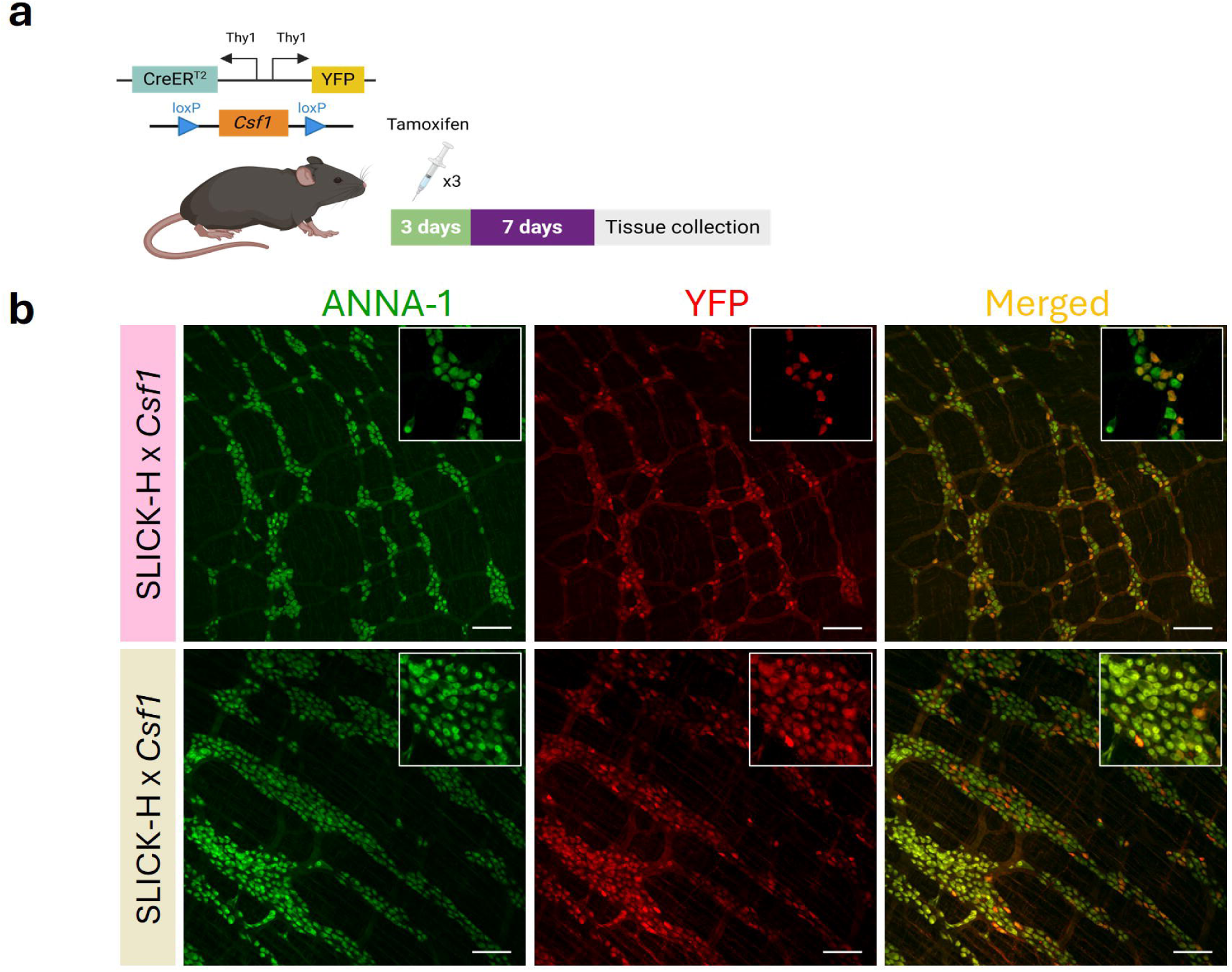
Validation of neuronal Csf1 deletion and an independent genetic model of neuron-derived CSF1. (a) Schematic of tamoxifen-induced recombination in SLICK-Hx*Csf1* mice. (b) Representative whole-mount images of ANNA-1⁺ enteric neurons and YFP^+^ cells in control and SLICK-Hx*Csf1* mice, confirming recombination within the enteric neuronal compartment.

## Author contributions

**KW**: Conceptualization of the work, performance of experiments, experimental supervision, data analysis, writing of the original draft, review of the original draft and editing. **JSOC**: Performance of experiments, review of the original draft and editing. **MD**: Performance of experiments, visualization, review of the original draft and editing. **BD**: Data curation and analysis, visualization, review of the original draft and editing. **YL**: Data curation and analysis, visualization, review of the original draft and editing. **SJ**: Performance of experiments, review of the original draft and editing. **MM**: Performance of experiments, review of the original draft and editing. **MHE**: Performance of experiments, review of the original draft and editing. **NW**: Performance of experiments, review of the original draft and editing. **XC**: Performance of experiments, review of the original draft and editing. **JXJ:** Intellectual input, review of the original draft and editing. **MG:** Intellectual input, review of the original draft and editing. **KAS:** Intellectual input, review of the original draft and editing. **AB**: Intellectual input, review of the original draft and editing. **GF**: Intellectual input, review of the original draft and editing. **GC**: Conceptualization of the work, provision of resources, writing of the original draft, study supervision, review of the original draft and editing.

## Acknowledgements

The authors thank the Mayo Clinic Flow Cytometry Core and Genome Analysis Core for their technical support, data acquisition, and expert guidance throughout this project.

## Data Sharing

All data generated or analyzed in this study are included in the article and its supplementary materials; additional information can be obtained from the corresponding authors upon request.

---

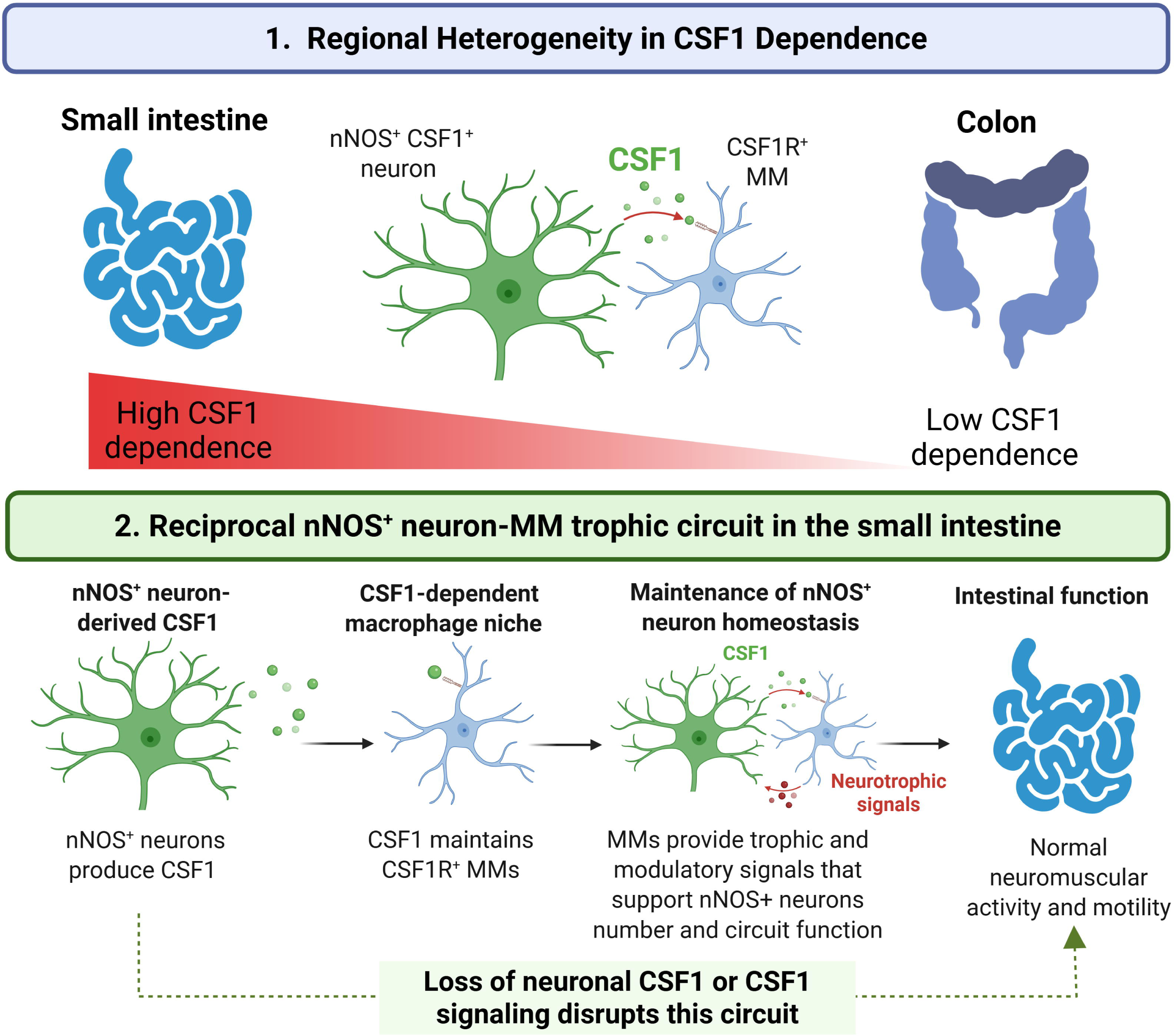

